# Influenza A virus-induced thymus atrophy differentially affects dynamics of conventional and regulatory T cell development

**DOI:** 10.1101/2020.09.09.274795

**Authors:** Yassin Elfaki, Philippe A. Robert, Christoph Binz, Christine S. Falk, Dunja Bruder, Immo Prinz, Stefan Floess, Michael Meyer-Hermann, Jochen Huehn

**Affiliations:** Department of Experimental Immunology, Helmholtz Centre for Infection Research, Braunschweig, Germany; Department of Systems Immunology and Braunschweig Integrated Centre of Systems Biology, Helmholtz Centre for Infection Research, Braunschweig, Germany; Institute of Immunology, Hannover Medical School, Hannover, Germany; Institute of Transplant Immunology, Hannover Medical School, Hannover, Germany; Infection Immunology Group, Institute of Medical Microbiology, Infection Control and Prevention, Health Campus Immunology, Infectiology and Inflammation, Otto-von-Guericke University Magdeburg, Magdeburg, Germany; Immune Regulation Group, Helmholtz Centre for Infection Research, Braunschweig, Germany; Cluster of Excellence RESIST (EXC 2155), Hannover Medical School, Hannover, Germany; Institute for Biochemistry, Biotechnology and Bioinformatics, Technical University Braunschweig, Braunschweig, Germany

**Author notes:** The author contributed equally to this work (Yassin Elfaki, Philippe A. Robert). correspondence to: Jochen Huehn, Department of Experimental Immunology, Helmholtz Centre for Infection Research, Inhoffenstr. 7, 38124 Braunschweig, Germany.

**Keywords:** thymus, Foxp3^+^, Treg cells, thymus atrophy, influenza A virus, mathematical modeling, ordinary differential equations

## Abstract

Foxp3^+^regulatory T (Treg) cells, which are crucial for maintenance of self-tolerance, mainly develop within the thymus, where they arise from CD25^+^Foxp3^-^or CD25^-^Foxp3^+^ Treg cell precursors. Although it is known that infections can cause transient thymic involution, the impact of infection-induced thymus atrophy on thymic Treg (tTreg) cell development is unknown. Here, we infected mice with influenza A virus (IAV) and studied thymocyte population dynamics post infection. IAV infection caused a massive, but transient thymic involution, dominated by a loss of CD4^+^CD8^+^ double-positive (DP) thymocytes, which was accompanied by a significant increase in the frequency of CD25^+^Foxp3^+^ tTreg cells. Differential apoptosis susceptibility could be experimentally excluded as a reason for the relative tTreg cell increase, and mathematical modeling suggested that enhanced tTreg cell generation cannot explain the increased frequency of tTreg cells. Yet, an increased death of DP thymocytes and augmented exit of single-positive (SP) thymocytes was suggested to be causative. Interestingly, IAV-induced thymus atrophy resulted in a significantly reduced T cell receptor (TCR) repertoire diversity of newly produced tTreg cells. Taken together, IAV-induced thymus atrophy is substantially altering the dynamics of major thymocyte populations, finally resulting in a relative increase of tTreg cells with an altered TCR repertoire.

## INTRODUCTION

CD4^+^ regulatory T (Treg) cells, which express the lineage-specification factor forkhead box P3 (Foxp3), constitute a subset of CD4^+^ T cells that is crucial for the maintenance of immune homeostasis and self-tolerance [1]. The majority of the Treg cell population (approximately 80 %) originates from the thymus, and hence these cells are termed thymus-derived Treg (tTreg) cells [2]. In the thymus, the development of tTreg cells proceeds through a two-step process involving T cell receptor (TCR) and cytokine signaling that first produces CD25^+^Foxp3^-^ or CD25^-^Foxp3^+^ Treg cell precursors, which in a second step contribute almost equally to the generation of mature CD25^+^Foxp3^+^ tTreg cells [3-6].

Thymus atrophy is a condition whereby the size and function of the thymus are greatly reduced. Many stimuli have been shown to result in thymus atrophy, including stress, inflammation, infections, malnutrition, pregnancy, and aging [7-12]. The evolutionary benefit of thymus atrophy is still debated, with both metabolic and immunologic hypotheses being proposed. For instance, infection-induced thymus atrophy is thought to be beneficial, since a temporary cessation in T cell production might limit the effect of dominant tolerance on the course of the infection [13]. Interferon-α has been reported as a critical molecular mediator of infection-induced thymus atrophy, and sensitivity of the thymic epithelium is controlled by a sophisticated microRNA network [14, 15]. It is important to note that thymic involution during infection is a very rapid yet transient phenomenon, with recovery within one to two weeks [14]. However, despite this ample knowledge it has not yet been elucidated whether infection-induced thymus atrophy leads to changes in tTreg cell development.

Mathematical modeling has been used to describe the dynamics of thymocyte populations at steady state [16] and following drug-induced atrophy [17, 18], or to unravel determinants of the CD4:CD8 ratio in which T cells are generated by thymopoiesis [19]. However, these models did not consider mature tTreg cells and their precursors, and a new model is needed to understand tTreg cell dynamics during infection-induced atrophy.

Influenza A virus (IAV) is a single-stranded RNA virus belonging to *Orthomyxoviridae* family of viruses. It is an important respiratory pathogen that caused many epidemics and pandemics [20]. IAV has been shown to induce thymus atrophy accompanied by lymphopenia, and this IAV-induced thymus atrophy is mainly characterized by loss of CD4^+^CD8^+^ double-positive (DP) thymocytes [21]. Several mechanisms have been suggested to contribute to IAV-induced thymus atrophy, including direct interference with T cell development, elevated IFNγ production by innate CD8α^+^ T cells and/or NK cells in the thymus, and increased levels of glucocorticoids [21-23]. Yet, consequences of IAV-induced thymus atrophy on tTreg cell development are unknown.

In the present study, we have investigated the dynamics of major thymocyte populations during IAV-induced thymus atrophy. The infection-induced thymic involution was dominated by a rapid yet transient loss of DP thymocytes and a simultaneous significant increase in the frequency of newly generated CD25^-^Foxp3^+^ precursors and CD25^+^Foxp3^+^ tTreg cells, the latter interestingly showing a significantly reduced TCR repertoire diversity. When studying the molecular mechanisms underlying this phenomenon, mathematical modeling suggested an involvement of an increased death of DP thymocytes and an augmented exit of SP thymocytes from the thymus to be causative for the relative increase in newly generated CD25^+^Foxp3^+^ tTreg cells at the peak of IAV-induced thymus atrophy.

## RESULTS

### Frequency of tTreg cells and their CD25^-^Foxp3^+^ precursors increases upon IAV-induced thymus atrophy

IAV infection has been reported to cause a transient thymic involution [21, 23]. In order to study the impact of IAV-induced thymus atrophy on tTreg cell development, we performed a kinetic analysis by infecting adult Foxp3^hCD2^xRag1^GFP^ double reporter mice with a mouse-adapted strain of IAV (PR8M) and analyzing their thymi at different time points post infection. Under the experimental conditions chosen, thymus atrophy was evident most dramatically 10 days post infection (dpi), both macroscopically and regarding total cellularity (Fig. 1A). In line with previously published data [23], this drop was attributable to a loss of DP thymocytes (Fig. 1B, Supporting Information Fig. 1 and ‘Supporting Information Mathematical Model’). Surprisingly, coinciding with the peak of atrophy, the frequency of CD25^+^Foxp3^hCD2+^ Treg cells among CD4 single-positive (SP) thymocytes increased significantly in thymi from IAV-infected mice when compared to PBS-treated controls (Fig. 1C and Supporting Information Fig. 1). Furthermore, a modest, though not significant, increase in the frequency of CD25^-^Foxp3^hCD2+^, but not CD25^+^Foxp3^hCD2-^ Treg cell precursors was observed (Fig. 1C and Supporting Information Fig. 1). Taken together, these data show that IAV-induced thymus atrophy results in an increase in the frequency of tTreg cells and their CD25^-^Foxp3^+^ precursors.

**Figure 1.**
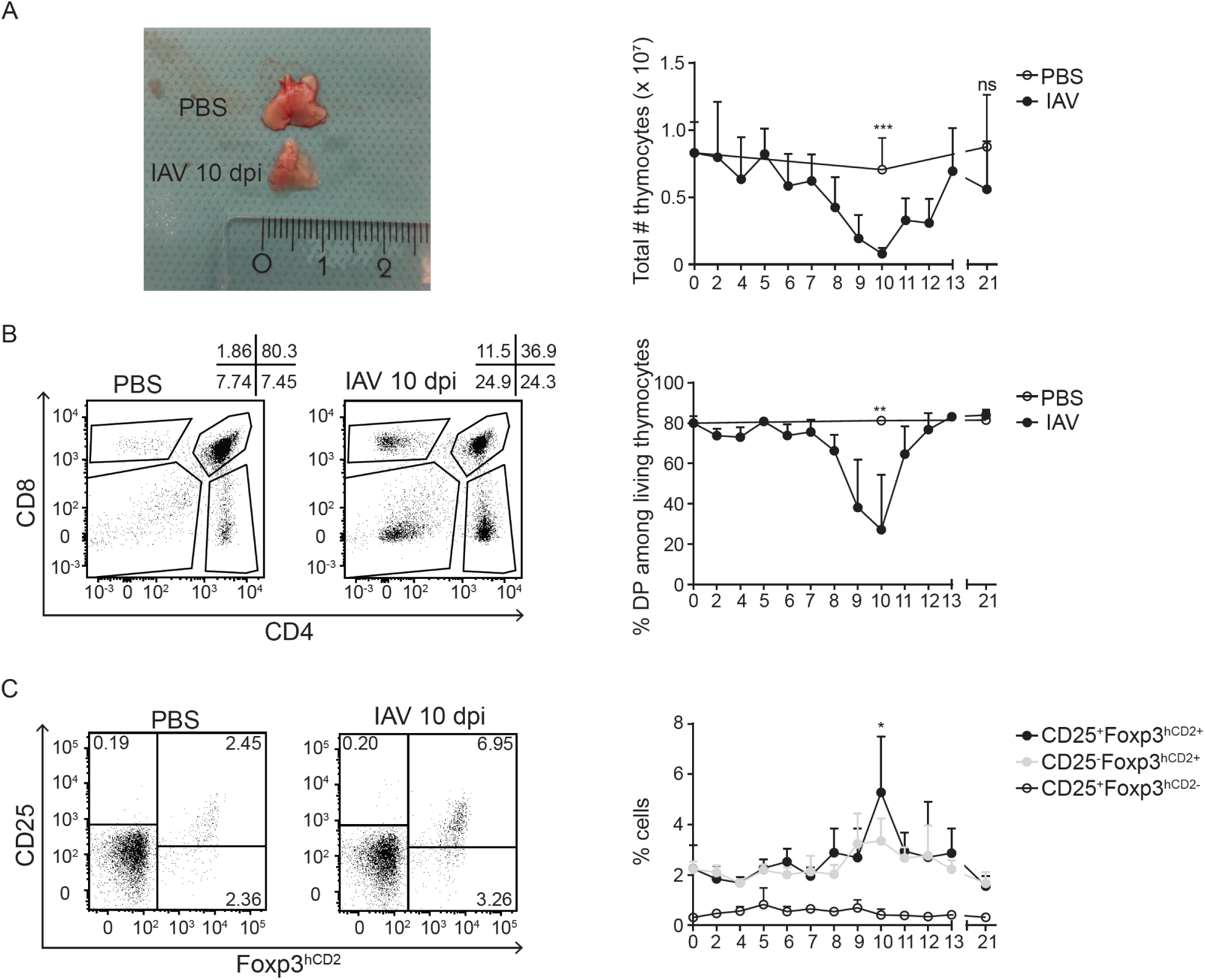
Frequency of tTreg cells and their CD25^-^Foxp3^+^ precursors increases upon IAV-induced thymus atrophy. Foxp3^hCD2^xRag1^GFP^ double reporter mice were infected with IAV and control mice received PBS. Thymi were analyzed at different days post infection (dpi). (A) (Left) Thymic morphology and gross size of PBS-treated and IAV-infected mice at 10 dpi. Representative of 9-20 animals per group. (Right) Graph summarizes thymic cellularity collected from PBS-treated (open circles) and IAV-infected (filled circles) mice at indicated dpi. (B) (Left) Representative dotplots depict gating strategy to identify CD4SP, CD8SP, DP and DN populations among living thymocytes prepared from PBS-treated and IAV-infected mice at 10 dpi. Numbers indicate frequencies of cells in the respective indicated gates. (Right) Graph summarizes frequency of DP thymocytes among living thymocytes prepared from PBS-treated (open circles) and IAV-infected (filled circles) mice at the indicated dpi. (C) (Left) Representative dotplots depict gating strategy to identify CD25^+^Foxp3^hCD2+^ Treg cells, CD25^-^Foxp3^hCD2+^ Treg cell precursors and CD25^+^Foxp3^hCD2-^ Treg cell precursors among CD4SP thymocytes from PBS-treated and IAV-infected mice at 10 dpi. Numbers indicate frequencies of cells in indicated gates. (Right) Graph summarizes frequencies of CD25^+^Foxp3^hCD2+^ Treg cells (filled circles) and their precursors (CD25^+^Foxp3^hCD2-^ (open circles) and CD25^-^Foxp3^hCD2+^ (grey circles)) among CD4SP thymocytes at the indicated dpi. Data were pooled from at least two independent experiments with 5-9 mice per group and presented as mean + SD. Each dot represents mean of biological replicates. Mann-Whitney test was used to test for statistical significance and significance was calculated by comparing CD25^+^Foxp3^hCD2+^ Treg cells, CD25^-^Foxp3^hCD2+^ Treg cell precursors and CD25^+^Foxp3^hCD2-^ Treg cell precursors from IAV-infected mice at 10 dpi and PBS-treated controls (not depicted).

### IAV infection results in increased frequency of newly generated tTreg cells

Treg cells in the thymus are a diverse population, consisting of newly generated *bona fide* tTreg cells as well as recirculating Treg cells, which re-enter the thymus from the periphery [24-29]. We next asked if the abovementioned overrepresentation of Treg cells within atrophied thymi maps to the newly generated or recirculating Treg cells, or both. Hence, we here made use of transgenic Rag1^GFP^ reporter mice, which express green fluorescent protein (GFP) under control of the recombination-activating gene 1 (*Rag1*) promoter [30]. In these mice, GFP expression identifies newly generated thymocytes and discriminates them from older and/or re-circulating T cells. In line with previously published data [24], the vast majority of CD4SP thymocytes was newly generated (Rag1^GFP+^) (Fig. 2A and Supporting Information Fig. 1). Interestingly, IAV-infected mice at 10 dpi showed a significant increase in the frequency of Rag1^GFP-^ CD4SP thymocytes when compared to PBS-treated controls, yet their absolute numbers remained unchanged (Fig. 2A and Supporting Information Fig. 2A).

**Figure 2.**
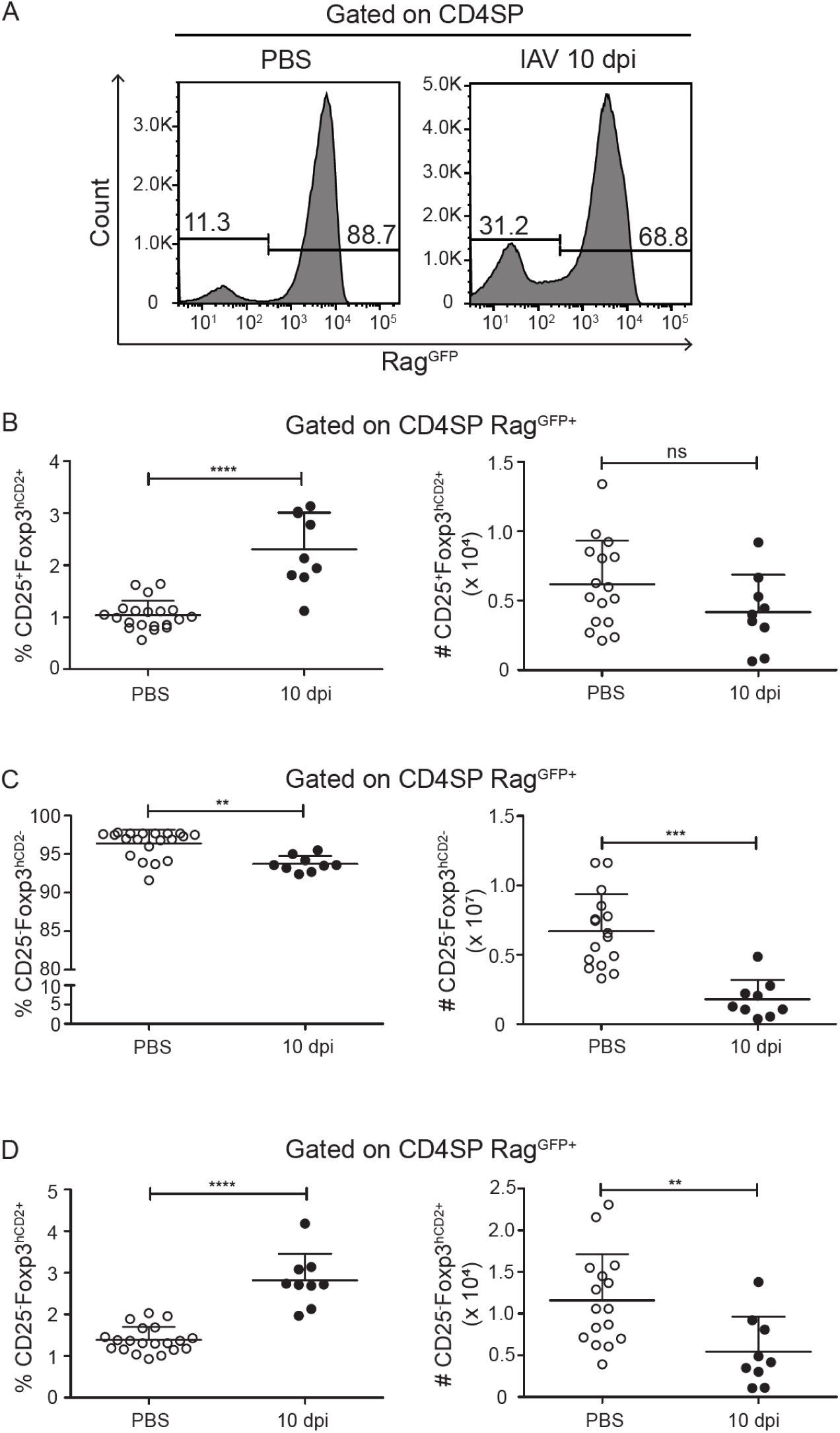
Frequency of *de novo* novo generated tTreg cells increases upon IAV infection. Foxp3^hCD2^xRag1^GFP^ double reporter mice were infected with IAV and control mice received PBS. Thymi were analyzed at different dpi. (A) Representative histograms depict Rag1^GFP^ expression among CD4SP thymocytes from PBS-treated and IAV-infected mice at 10 dpi. Numbers indicate frequencies of cells in indicated gates. (B-D) Scatterplots summarize frequencies (left) and absolute numbers (right) of CD25^+^Foxp3^hCD2+^ Treg cells (B), CD25^-^Foxp3^hCD2-^ Tconv (C) and CD25^-^Foxp3^hCD2+^ Treg cell precursors (D) among CD4SP Rag1^GFP+^ cells from PBS-treated (open circles) and IAV-infected (filled circles) mice at 10 dpi. Data were pooled from three independent experiments with 9-20 mice per group and presented as mean + SD. Each dot represents an individual mouse. Mann-Whitney test was used to test for statistical significance.

Among newly generated Rag1^GFP+^ CD4SP thymocytes, we observed a significant increase in the frequency of CD25^+^Foxp3^hCD2+^ Treg cells in IAV-infected mice 10 dpi when compared to PBS-treated controls (Fig. 2B). Conversely, the frequency of CD25^-^Foxp3^hCD2-^ Tconv decreased significantly in thymi from IAV-infected mice compared to those from control mice (Fig. 2C). Surprisingly, unlike the numbers of Tconv, which were significantly reduced in atrophied thymi (Fig. 2C), the numbers of Treg cells were only slightly decreased, not reaching statistical significance (Fig. 2B). In addition, in thymi from IAV-infected mice at 10 dpi the frequency of CD25^-^Foxp3^hCD2+^ Treg cell precursors increased significantly compared to those from PBS-treated control mice, suggesting favored differentiation of Treg cells via these precursors in this setting (Fig. 2D).

When we tested the Treg cell and Tconv populations among recirculating Rag1^GFP-^ CD4SP thymocytes, we neither observed an increase in frequencies nor numbers when thymi from IAV-infected mice 10 dpi were compared to those from PBS-treated controls (Supporting Information Fig. 2B+C), indicating that the increase in Treg cell frequency seen in thymi of IAV-infected mice does not result from an increased recirculation of Treg cells during infection-induced thymus atrophy.

### Intrathymic Treg cells and Tconv show comparable survival rates during IAV-induced thymus atrophy

The data from the kinetic analysis indicate that newly generated tTreg cells are favored over their Tconv counterparts upon IAV-induced thymus atrophy. To get insights into the underlying molecular mechanism we first tested whether the relative increase of tTreg cells is a result of a superior ability to survive when compared to Tconv in the context of IAV-induced thymus atrophy by staining for active caspase 3/7. Examining the newly generated Rag1^GFP+^ CD4SP thymocytes revealed minimal propensity to apoptosis among both tTreg cells and Tconv as hardly any caspase 3/7^+^ apoptotic cells could be detected (Fig. 3A). Interestingly, no impact of infection-induced thymus atrophy on survival of newly generated Treg cell and Tconv was observed at 7, 10 and 14 dpi, and both cell types showed similarly low frequencies of caspase 3/7^+^ cells at all time points (Fig. 3B). In contrast, many more cells were undergoing apoptosis among recirculating Rag1^GFP-^ CD4SP thymocytes, yet again no impact of infection-induced thymus atrophy and no differences between Treg cell and Tconv were observed (Fig. 3C+D). Together, these data indicate that the relative enrichment of newly generated tTreg cells is unlikely to result from a better survival compared to newly generated Tconv.

**Figure 3.**
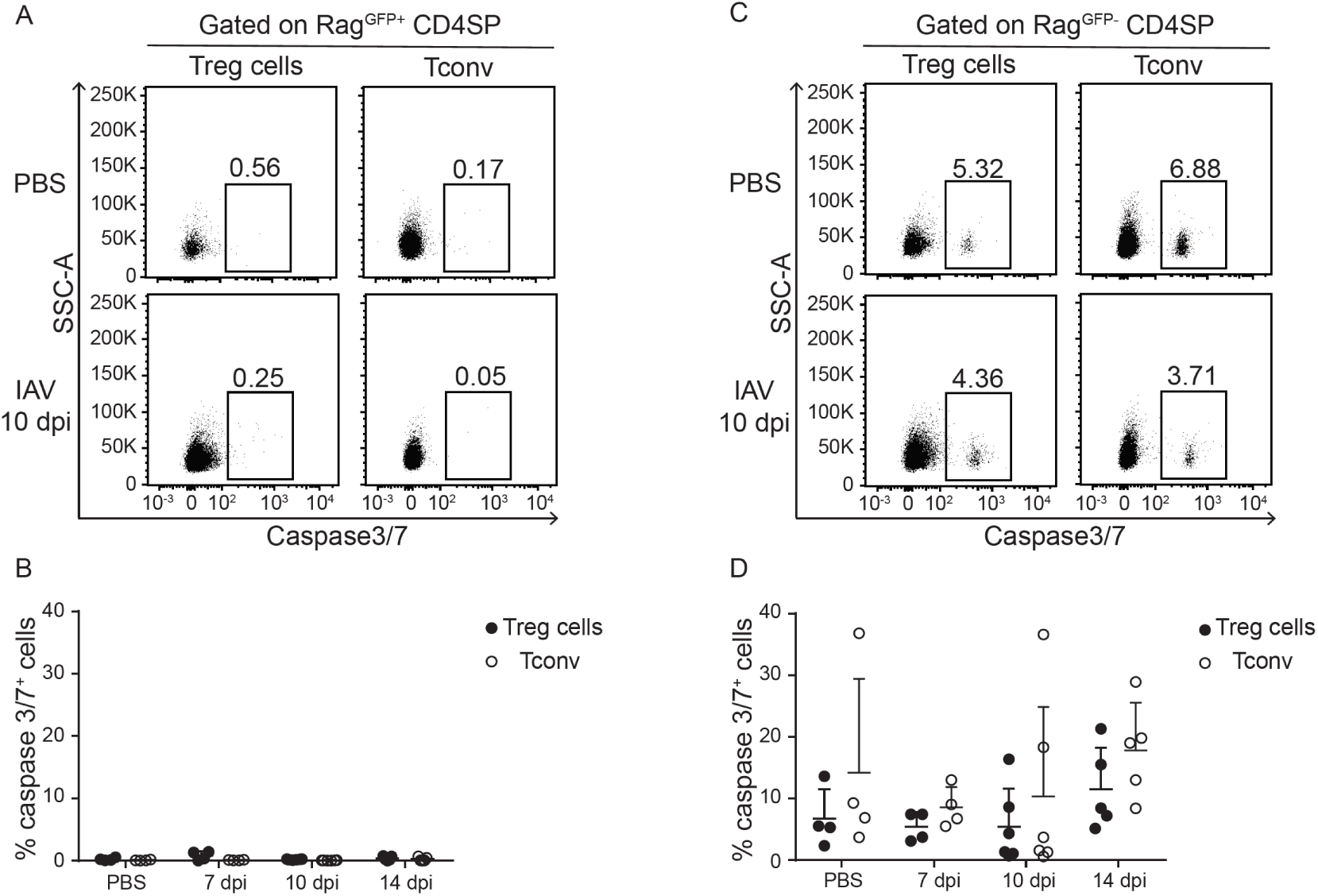
Intrathymic Tregs and Tconv show comparable survival rates during IAV-induced thymic atrophy. Foxp3^hCD2^Rag1^GFP^ double reporter mice were infected with IAV and control mice received PBS. Thymi were analyzed at different dpi. (A) Representative dotplots depict caspase 3/7^+^ cells among Treg cells and Tconv pregated on CD4SP Rag1^GFP+^ cells from thymi of PBS-treated and IAV-infected mice at 10 dpi. Numbers indicate fractions of cells in indicated gates. (B) Scatterplots summarize caspase 3/7^+^ cells among Treg cells (filled circles) and Tconv (open circles) pregated on CD4SP Rag1^GFP+^ cells. (C) Representative dotplots depict caspase 3/7^+^ cells among Treg cells and Tconv gated on CD4SP Rag1^GFP-^ cells from thymi of PBS-treated and IAV-infected mice at 10 dpi. Numbers indicate fractions of cells in indicated gates. (D) Scatterplots summarize caspase 3/7^+^ cells among Treg cells (filled circles) and Tconv (open circles) pregated on CD4SP Rag1^GFP-^ cells. Data were pooled from two independent experiments with 4-8 mice per group and presented as mean + SD. Each dot represents an individual mouse. Mann-Whitney test was used to test for statistical significance.

### Newly generated CD4SP thymocytes show unaltered TCR signaling upon IAV-induced thymus atrophy

TCR signaling is critical for tTreg cell development [3, 31], and reduced TCR signaling has been shown to enhance Treg cell differentiation in multiple models [32-34]. To test whether IAV-induced thymus atrophy is accompanied by reduced TCR signaling, we sorted newly generated Rag1^GFP+^ CD4SP thymocytes from IAV-infected mice at 9 dpi, shortly before the peak of thymus atrophy, and stained them for CD5 and Nur77, the levels of which correlate with the intensity of TCR signaling and negative selection [35, 36]. Flow cytometric analysis revealed that Rag1^GFP+^ CD4SP thymocytes from IAV-infected mice showed similar mean fluorescence intensity levels for both CD5 and Nur77 when compared to newly generated Rag1^GFP+^ CD4SP thymocytes taken from PBS-treated control mice (Fig. 4A+B). Hence, these data suggest that a reduced TCR signaling is unlikely to be causative for the relative tTreg cell enrichment upon IAV-induced thymus atrophy.

**Figure 4.**
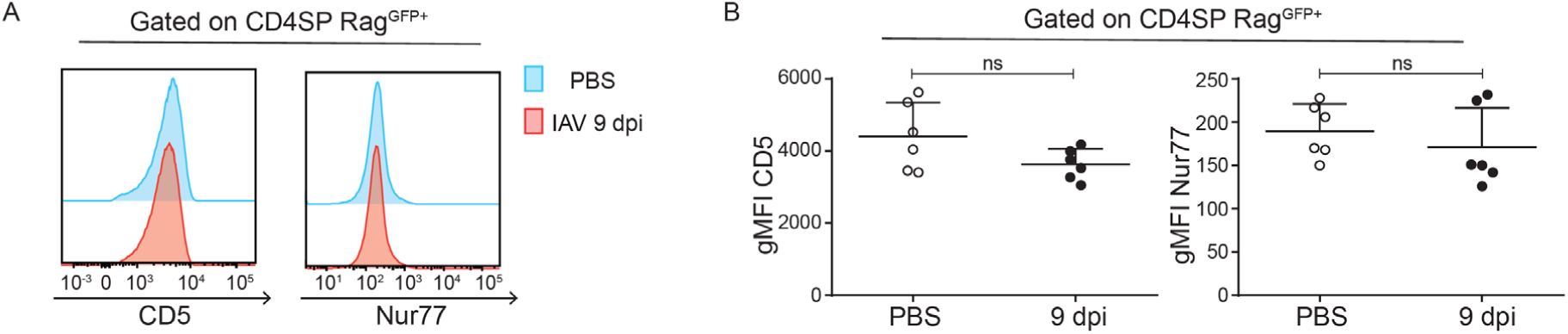
*De novo* generated CD4SP thymocytes show comparable levels of CD5 and Nur77 upon IAV-induced thymus atrophy. Foxp3^hCD2^xRag1^GFP^ double reporter mice were infected with IAV and control mice received PBS. Thymi were collected, and CD4SP Rag1^GFP+^ cells were sorted and analyzed. (A) Representative histograms display CD5 (left) and Nur77 (right) expression in sorted CD4SP Rag1^GFP+^ cells from thymi of PBS-treated (blue) and IAV-infected (red) mice at 9 dpi. (B) Scatterplots summarize geometric mean fluorescence intensity (gMFI) of CD5 (left) and Nur77 (right) in sorted CD4SP Rag1^GFP+^ cells from thymi of PBS-treated and IAV-infected mice at 9 dpi. Data were pooled from two independent experiments which included six mice per group and presented as mean + SD. Each dot represents an individual mouse. Mann-Whitney test was used to test for statistical significance.

### Intrathymic levels of TGF-β transiently increase upon IAV-induced thymus atrophy

In addition to signals provided via the TCR also cytokines are essential mediators for Treg cell development in the thymus [37]. Thus, in order to assess the role of cytokines in the preferential increase of tTreg cells in the atrophied thymus, we measured their levels at different time points of IAV-induced thymus atrophy. Levels of IL-2, a key cytokine for tTreg cell development, were unchanged at 7, 9 and 12 dpi when compared to PBS-treated controls (Fig. 5A). Similarly, levels for the inflammatory cytokines IL-1α, IL-β and IFNγ, all known to modulate tTreg cell development or be involved in thymus atrophy [22, 28], did not increase in the thymus upon infection-induced atrophy. Only slightly reduced IL-6 levels were observed in thymi at 12 dpi when compared to PBS-treated controls (Supporting Information Fig. 3). Interestingly, levels of TGF-β, for which a positive effect on tTreg cell development was reported [38], were significantly elevated during infection-induced thymus atrophy at 7 dpi, before returning to levels of PBS-treated controls at 12 dpi (Fig. 5B). These data suggest that increased levels of TGF-β might contribute to an enhanced tTreg cell differentiation during IAV-induced thymus atrophy.

**Figure 5.**
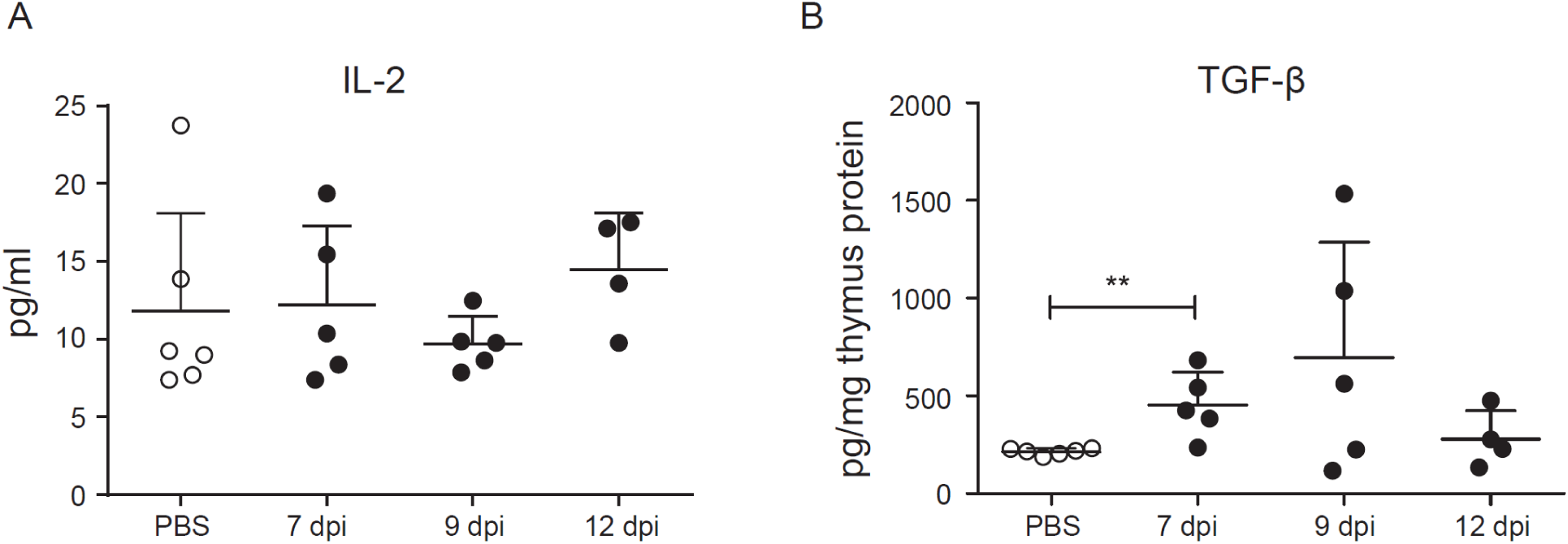
TGF-β but not IL-2 in thymus increases significantly upon thymus atrophy. Foxp3^hCD2^xRag1^GFP^ double reporter mice were infected with IAV and control mice received PBS. Thymi were analyzed at indicated dpi. Concentrations of IL-2 and TGF-β were measured by Bio-plex 23-plex mouse kit or LEGENDplex kit, respectively. Scatterplots summarize IL-2 (A) and TGF-β (B) concentrations in thymi from PBS-treated (open circles) and IAV-infected mice (filled circles) at indicated dpi. Data were pooled from two independent experiments with 4-6 mice per group and presented as mean + SD. Each dot represents an individual mouse. Mann-Whitney test was used to test for statistical significance.

### Mathematical modeling suggests increased DP thymocyte death and augmented exit of SP thymocytes as cause for the increased frequency of tTreg cells upon IAV-induced thymus atrophy

Next, we analyzed possible mechanisms underlying the relative enrichment of newly generated tTregs upon IAV-induced thymus atrophy with a mathematical model simulating the time evolution of thymocyte populations during IAV infection, including Treg cells and their precursors, using the data from the kinetic study (Fig. 6A). Each node represents a population and arrows represent the mechanisms of differentiation, proliferation or death (for details see ‘Design of the mathematical model’ in the Material and Methods section). The DP thymocyte compartment is built on a previous model of T cell development and population dynamics, where the early proliferating DP (eDP) and late, resting DP thymocytes (lDP) are distinguished [17]. We added the following Rag1^+^ CD4SP thymocyte populations: CD25^-^Foxp3^-^ cells (Tconv), CD25^+^Foxp3^-^ Treg cell precursors, CD25^-^Foxp3^+^ Treg cell precursors, and CD25^+^Foxp3^+^ mature Treg cells. Furthermore, as several studies have highlighted that the two Treg cell precursor populations arise at different time points of T cell development and harbor differential dynamics [4-6, 27], we compared different model structures for Treg cell development (Supporting Information Fig. 4), namely where both Treg cell precursors either arise from DP thymocytes (structure A) or Tconv (structure B), or where CD25^-^Foxp3^+^ Treg cell precursors arise from Tconv, while CD25^+^Foxp3^-^ Treg cell precursors directly arise from DP thymocytes (structure C).

**Figure 6.**
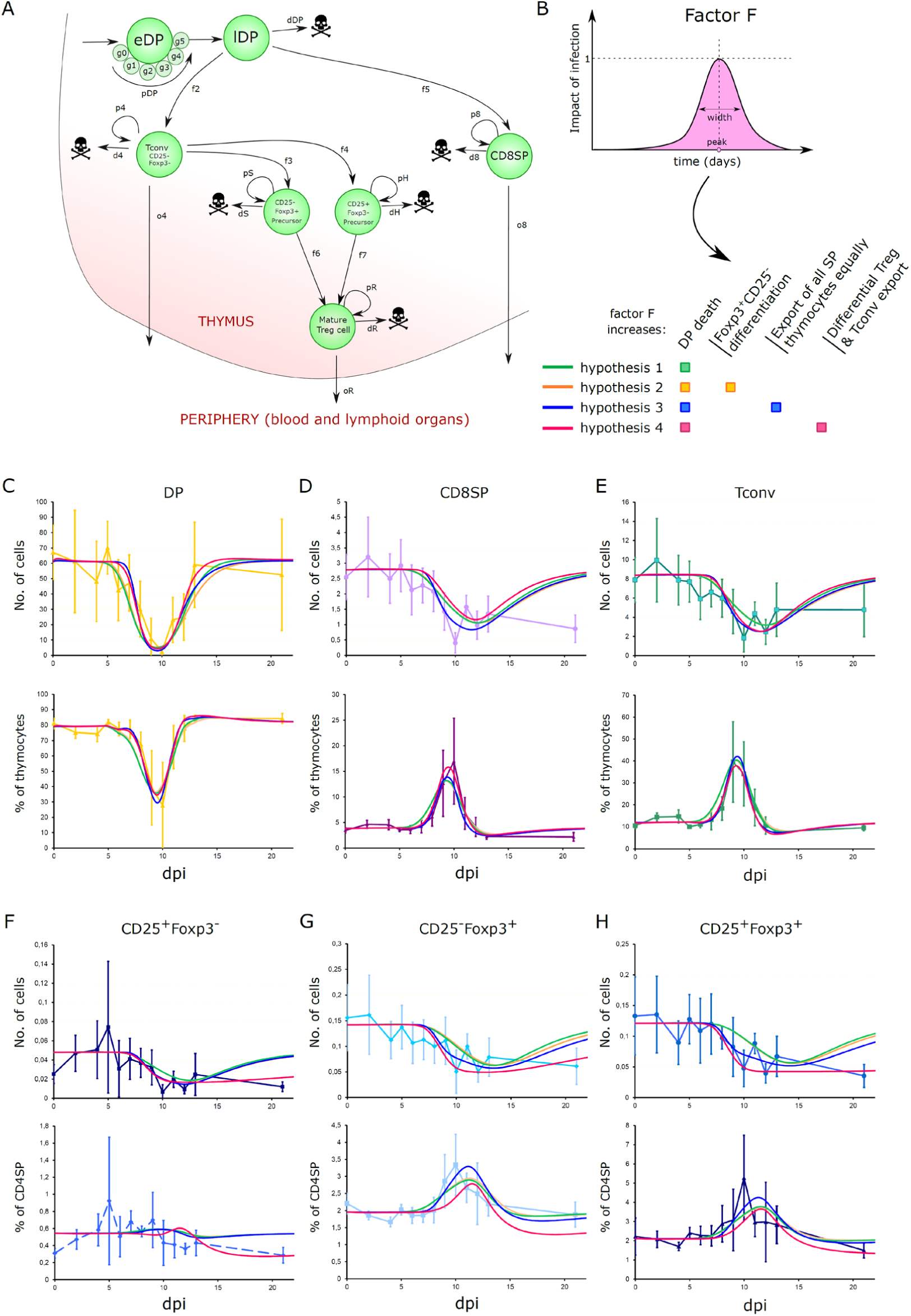
Dynamical modeling of thymus kinetics under IAV-induced atrophy under different hypotheses. (A) Mathematical model for DP thymocyte, Tconv, CD8SP thymocyte, Treg cell precursors and mature Treg cell compartments. (B) (Top) Factor F, used to model the effect of IAV infection on the thymus. (Bottom) Different hypotheses on the effect of IAV infection. (C-H) Best fitted curves to explain experimental kinetics of all subpopulations with each hypothesis, as absolute numbers (top) and frequencies (bottom). The colors of the curves match the hypotheses in B. Absolute numbers are only shown for the newly generated Rag1^GFP+^ populations. Frequencies show the summed percentage of Rag1^GFP+^ and Rag1^GFP-^ cells together.

The effect of IAV infection is modeled as a perturbating factor F, with a Gaussian dynamic over time (Fig. 6B), whose peak time and width are to be found. To discriminate possible mechanisms responsible for the dynamics of the thymocyte populations during IAV-induced thymus atrophy, we tested different assumptions, namely Factor F increases DP thymocyte death, Factor F increases *de novo* CD25^-^Foxp3^+^ Treg cell precursor generation, Factor F increases the export of all SP thymocyte populations equally, or Factor F increases the export of Tconv compared to Treg cells and Treg cell precursors differentially (Fig. 6B). Deliberately, we did not test an increased cell death among Tconv or Treg cell precursors, since we did not detect any significant cell death among newly generated CD4SP thymocyte subsets (Fig. 3). Fitting the data under these assumptions identified the increase in DP thymocyte death as a dominant factor that could already explain for the most data the dynamics of all major thymocyte populations, including the transient increase in the frequency of mature tTreg cells and their CD25^-^Foxp3^+^ precursors during the peak of the IAV-induced thymus atrophy (green curve in Fig. 6C-H and Supporting Information Fig. 5-7). If other assumptions like an increased *de novo* CD25^-^Foxp3^+^ Treg cell precursor generation or an increased export of all SP thymocyte populations are simulated alone, the DP thymocyte shrinkage dynamics could not be explained, as can be seen by the very high costs (Supporting Information Fig. 8).

While increased DP thymocyte death was compatible with most data for the three model structures, the peaks of the fractions of mature tTreg cells and their CD25^-^Foxp3^+^ precursors appeared earlier and were slightly higher in the experimental measurement when compared to the simulations (green curve in Fig. 6C-H and Supporting Information Fig. 5-7). Thus, we tested combinations of assumptions.

At first, we simulated atrophy under both increased DP thymocyte death and increased *de novo* generation of CD25^-^Foxp3^+^ Treg cell precursors (hypothesis 2 in Fig. 6C-H, yellow curves). However, the best parameter values (see Supporting Information Fig. 9 and ‘Design of the mathematical model’ in the Material and Methods section) suggests an only marginally 1.00006-fold increased *de novo* generation of CD25^-^Foxp3^+^ Treg cell precursors (yellow curve in Fig. 6 partially overlapping with the green curve, and Supporting Information Fig. 5+6). A sensitivity analysis around the best curve showed that a substantially modulated *de novo* generation of CD25^-^Foxp3^+^ Treg cell precursors would be detectable due to different dynamics of the Treg cell precursors (Supporting Information Fig. 10A). The corresponding identifiability analysis showed that the increased *de novo* generation of CD25^-^Foxp3^+^ Treg cell precursors also increases the cost of the fittings as it has a strong impact on the dynamics of absolute cell numbers (Supporting Information Fig. 10B). Thus, increased *de novo* generation of CD25^-^Foxp3^+^ Treg cell precursors does not provide a better fit nor explanation of the dataset (Supporting Information Fig. 9).

We next tested an increased export of all SP thymocytes (CD8SP thymocytes, Tconv, Treg cells and both Treg cell precursors) with equal strength and simulated this together with the assumption of an increased DP thymocyte death (hypothesis 3 in Fig. 6B). This is supported by the observation that the number of CD8SP thymocytes and Tconv seems to decay before the predicted effect of the increased DP thymocyte death (green curve in Fig. 6D+E). The best fit improved the dynamics of CD8SP thymocytes and Tconv, and also the transient increase in the frequency of mature tTreg cells and their CD25^-^Foxp3^+^ precursors was refined (blue curves in Fig. 6D+E+G+H and Supporting Information Fig. 5+6). In general, the cost of each model structure was improved when compared to the assumption of an increased DP thymocyte death alone, and the best fit with model structure B had a significantly better corrected Akaike Information Criterion (AICc) when compared to the other model structures (Supporting Information Fig. 9).

Finally, we assumed a differential increase of the export of Tconv in comparison to Treg cells and their precursors, and simulated this together with increased DP thymocyte death (hypothesis 4). Although the cost of the best parameter set for this highly intuitive hypothesis was lower, indicating a better quantitative fitting (Supporting Information Fig. 9), the dynamics of the mature tTreg cells and their CD25^-^Foxp3^+^ precursors around the peak of the IAV-induced thymus atrophy were less well explained (red curves in Fig. 6G+H and Supporting Information Fig. 5+6). This discrepancy comes from the latest time point, where hypothesis 4 fits best, particularly regarding the absolute cell numbers, at the expense of a less good fit around the peak of the IAV-induced thymus atrophy.

For each model structure, hypotheses 3 and 4 are most well suited to describe the dynamics of all thymocyte populations during IAV-induced thymus atrophy (Supporting Information Fig. 9). While in model structure A and C, a differential export of Tconv and Tregs (hypothesis 4) does not impove the AICc index, model structure B shows an improvement of the AICc in this case. In order to discriminate between hypotheses 3 and 4, we predicted the half-life of both Treg cell precursor populations from the obtained best parameter sets, as an independent qualitative validation. In line with published data [5], hypothesis 3 within model structure B predicted realistic values of 0.5, 3.1 and 3.8 days half-life for CD25^+^Foxp3^-^precursors, CD25^-^Foxp3^+^ precursors and mature Treg cells, respectively, while hypothesis 4 generated values out of range (Supporting Information Fig. 9). We therefore rejected hypothesis 4 and kept model structure B with hypothesis 3 as the most likely scenario. In conclusion, the assumption of an increased DP thymocyte death is the major determinant for the dynamics of the thymocyte populations during IAV-induced atrophy, together with the assumption of an increased export of all SP thymocyte populations with equal strength. Furthermore, model structure B, where both Treg cell precursors arise from Tconv, is the best model structure in terms of cost, AICc values, and realistic half-lives of thymocyte subsets.

### IAV-induced thymus atrophy results in reduced TCR repertoire diversity among newly produced thymocytes

We next asked whether the IAV-induced thymus atrophy also affects the TCR repertoires of newly produced Treg cells and Tconv. To this end, CD25^+^Foxp3^hCD2+^ Treg cells and CD25^-^Foxp3^hCD2-^Tconv were isolated from both newly produced Rag1^GFP+^ as well as recirculating Rag1^GFP-^ CD4SP thymocytes of PBS-treated or IAV-infected mice 10 dpi. RNA was isolated from sorted cells and subjected to high-throughput next-generation sequencing of *Tcrb* transcripts to assess the TCR repertoire. Remarkably, the TCR repertoire diversity of the newly produced Treg cells was significantly reduced upon IAV infection (Fig. 7A) and was accompanied by a reduction of the rare clones (Fig. 7B). A similar pattern emerged when observing the newly produced Tconv, although not reaching statistical significance (Fig. 7A+B). Yet, the impact of IAV infection on the TCR repertoire diversity was less pronounced among the recirculating Treg cells and Tconv (Fig. 7A+B). Together, these data suggest that IAV-induced thymus atrophy causes a reduction in the TCR repertoire diversity of newly produced CD4SP thymocyte subsets, while having much less influence on the TCR repertoire of the recirculating T cell subsets.

**Figure 7.**
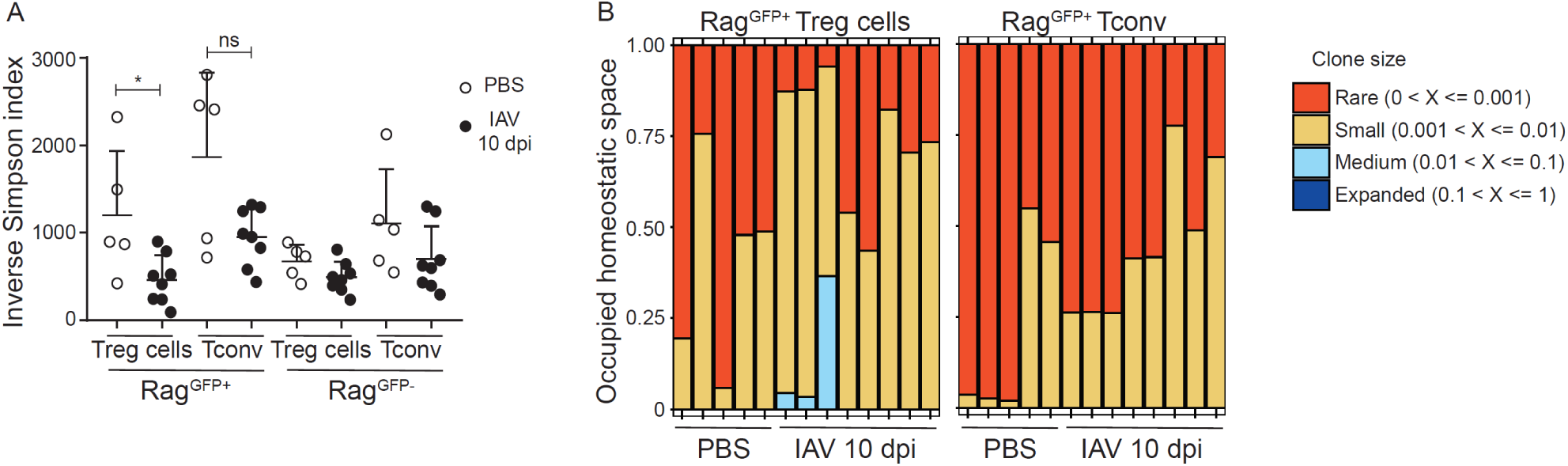
IAV-induced thymus atrophy results in reduced TCR repertoire diversity among *de novo* generated T cells. Foxp3^hCD2^xRag1^GFP^ double reporter mice were infected with IAV and control mice received PBS. Thymi were collected at 10 dpi, Rag1^GFP+^CD25^+^Foxp3^hCD2+^, Rag1^GFP+^CD25^-^Foxp3^hCD2-^, Rag1^GFP-^ CD25^+^Foxp3^hCD2+^, and Rag1^GFP-^CD25^-^Foxp3^hCD2-^ subsets of CD4SP thymocytes were sorted, and RNA from these cells was subjected to next-generation sequencing (NGS) of *Tcrb* transcripts. (A) Scatterplot summarizes the inverse Simpson index of thymic Rag1^GFP+^ Treg cells, Rag1^GFP+^ Tconv, Rag1^GFP-^ Treg cells and Rag1^GFP-^ Tconv isolated from either PBS-treated (open circles) or IAV-infected (filled circles) mice at 10 dpi. Each dot represents an individual mouse. (B) Cumulative bar graphs depict clonal homeostasis of Rag1^GFP+^ Treg cells and Rag1^GFP+^ Tconv sorted from thymi of PBS-treated and IAV-infected mice at 10 dpi. Each bar represents an individual mouse. Data were pooled from two independent experiments with 5-8 mice per group and presented as mean + SD. Mann-Whitney test was used to test for statistical significance.

## DISCUSSION

In the present study, we have shown that the massive, yet transient IAV-induced thymus atrophy is accompanied by transiently increased frequencies of mature tTreg cells and their CD25^-^Foxp3^+^precursors. This increase mapped to the newly generated Rag1^GFP+^Treg cells, and interestingly, while Rag1^GFP+^Tconv were significantly reduced in terms of both proportion and numbers, numbers of Rag1^GFP+^Treg cells were largely unaffected. Nonetheless, the relative increase of Treg cells was not the result of a better survival, as both Rag1^GFP+^ Treg cells and Tconv demonstrated similar (and negligible) propensity to apoptosis following IAV-induced thymus atrophy. Yet, the TCR repertoire diversity of both newly produced tTreg cells and Tconv was reduced upon IAV-induced thymus atrophy. Mathematical modeling suggested that the infection-induced increased death of DP thymocytes is the major determinant for the dynamics of the thymocyte populations during IAV-induced atrophy. It further proposed that an augmented exit of SP thymocytes could be involved in better describing the dynamics, rather than an enhanced tTreg cell generation being responsible for the relative increase in tTreg cells upon IAV-induced thymus atrophy. Our data argue for a model in which IAV-induced thymus atrophy differentially affects the dynamics of tTreg cells and Tconv.

Several previous reports had described the propensity of IAV infection to induce thymus atrophy [21-23]. Although different mechanisms have been suggested to contribute to thymus atrophy induced by IAV infection, such as IFNγ produced by NK cells or innate CD8α^+^ T cells, they all cause a loss of DP thymocytes, which is thought to result from an increase in serum corticosteroids [8]. Consistent with our data, a rapid rebound occurs and the thymus regains its usual shape and cellularity. Interestingly, IFNγ levels in thymi were not elevated upon infection in our experimental model, suggesting that IFNγ might not be essential in IAV-induced thymus atrophy. Since Liu et al. and Duan et al. injected IFNγ-neutralizing antibodies intraperitoneally [22, 23], it is possible that this neutralization attenuated the degree of inflammation, which subsequently reduced the severity of thymus atrophy. It remains to be defined whether intrathymic injection of IFNγ would have a similar alleviating effect on thymus atrophy.

Although thymic cellularity in the context of thymus atrophy was significantly decreased, the number of tTreg cells did not significantly change. In contrast, for Tconv both the frequency and absolute number were significantly reduced. This is thought to be deleterious to the host, since it is usually accompanied by lymphopenia [21]. In the context of IAV-induced thymus atrophy, the thymus may actively spare Treg cells in order to dampen the immune response after the peak of infection has passed. However, it is also possible that Treg cells are kept under tight control because their decrease, even in a transient manner, might result in the occurance of autoimmune episodes.

The finding that tTreg cells increase relative to Tconv upon IAV-induced thymus atrophy may be explained by various mechanisms, including a better survival of Treg cells over their Tconv counterparts. Yet, we could experimentally exclude this possibility by staining thymocytes for caspase 3/7. Interestingly, we saw a negligible fraction of cells undergoing apoptosis upon gating on Rag1^GFP+^cells, which represent the newly generated thymocytes, and furthermore, the frequency of caspase 3/7^+^ Treg cells was similar to that of Tconv. Intriguingly, apoptosis was most evident among the recirculating (Rag1^GFP-^) T cell subsets, and it will be very interesting to unravel the molecular mechanism underlying this differential susceptibility towards apoptosis between newly generated thymocytes and recirculating T cells in future studies.

The finding that tTreg cell generation is enhanced in the context of thymus atrophy has been shown before in a model of accelerated thymus atrophy [33]. In this model, accelerated thymus atrophy resulting from the induced deletion of FoxN1 was accompanied by disruption of negative selection [39]. The authors argue that the atrophied thymus attempts to balance this defect by actively generating tTreg cells. Indeed, they propose the decreased TCR signaling described to occur in this model to be the reason for this enhanced generation of tTreg cells [33]. Yet, in our model, the intensity of CD5 and Nur77 expression remained unchanged upon thymus atrophy. Furthermore, we had observed a significant increase in the frequency of only CD25^-^Foxp3^+^, but not CD25^+^Foxp3^-^ Treg cell precursors, and Farrar and colleagues have recently demonstrated that CD25^-^Foxp3^+^ precursors rely on a high TCR signal strength for their differentiation [6]. Thus, it is highly unlikely that a reduced TCR signaling is causative for the relative tTreg cell enrichment upon IAV-induced thymus atrophy.

The role of TGF-β in tTreg cell development is still a matter of debate. Initially, TGF-β signaling in the thymus was thought to be indispensable [38]. However, other reports claimed otherwise [40] or that it was only required for the survival of tTreg cells [41]. In 2014, Konkel *et al*. showed that apoptosis in the thymus is driving tTreg cell generation through TGF-β [42]. Since infection-induced thymus atrophy is accompanied by apoptosis of thymocytes, especially DP thymocytes [8], we asked whether TGF-β levels are increased in the context of IAV-induced thymus atrophy. Indeed, TGF-β levels increased significantly upon thymus atrophy. However, so far experimental data demonstrating a contribution of these increased TGF-β levels to an enhanced tTreg cell generation during IAV-induced thymus atrophy are missing. Furthermore, mathematical modeling did not provide any evidence for an involvement of this mechanism in the relative increase in tTreg cells upon IAV-induced thymus atrophy. Instead, the model suggested that the infection-induced increased death of DP thymocytes is the main driving factor, plus a possible contribution of an augmented exit of all SP thymocyte populations.

To the best of our knowledge, so far no mathematical model has accounted for the dynamic development of Treg cell precursors in the thymus. Despite the fact that a lot of progress has been made regarding our understating of tTreg cell development during the past years [2-6, 24-29], it is still not completely understood at which stage of T cell development the two Treg cell precursor populations arise, and it has been suggested that even earlier pre-precursors do exist [43]. Here, we show that the dynamics upon thymus atrophy contain information on Treg cell precursor dynamics. Further work would need to include quantitative datasets on the Treg cell populations’ properties, such as half-lives, BrdU labeling, or intra-thymic injection of Treg cell precursors, to experimentally validate the model. Further, although all model structures concord to delineate DP thymocyte death as the major determinant for the dynamics of the thymocyte populations during IAV-induced atrophy, the mechanistic impact of SP thymocyte export was leading to a better fit, judged by cost and AICc values. Since model structure B is the structure with the best fitting cost, our mathematical model is providing a first hint that Treg cell precursors might arise from CD4SP rather than directly from DP thymocytes.

The finding that CD25^-^Foxp3^+^ and CD25^+^Foxp3^-^ Treg cell precursors have different dynamics of Foxp3 induction [4-6, 27] might indicate that they recognize different types of self-antigens. We have previously shown that thymic selection can be simulated at single-cell level on the basis of time-integrated signaling to discriminate between Tconv and Treg cell differentiation [44]. It would be interesting to investigate if this thymic selection model would predict the diversity of tTreg cells and their precursors during infection-induced thymus atrophy.

The data of the present study demonstrate that IAV-induced thymus atrophy results in a decreased diversity of the TCR repertoire only among the newly generated T cells, both Treg cells and Tconv. Recirculating T cells were only slightly affected in terms of their TCR repertoire diversity. These data further support the notion that IAV-induced thymus atrophy is primarily affecting the process of T cell development and has only limited impact on recirculating T cells during and shortly after the infection. The narrowing of the TCR repertoire diversity could be a direct consequence of DP thymocyte death or the result of a functional impairment of thymic antigen-presenting cells (APCs) or simply a decrease in their numbers secondary to IAV-induced thymic atrophy. Thymic APCs are well known to critically contribute to tTreg cell development [45-49], and future studies need to unravel if IAV-induced thymus atrophy results in alterations of the thymic dendritic cell and thymic epithelial cell compartment, which finally affect the TCR repertoire diversity of newly generated T cell subsets. Indeed, it was shown in models of accelerated thymus atrophy and thymus atrophy due to aging that the functionality of thymic APCs was negatively influenced, resulting in a reduced ability to present self-antigens, which perturbed negative selection and finally provoked autoimmune phenotypes [50].

In summary, we have shown that IAV-induced thymus atrophy is strongly affecting the dynamics of major thymocyte populations, finally resulting in a relative increase of tTreg cells with an altered TCR repertoire. Our findings are particularly relevant given the frequent occurrence of thymus atrophy in different settings. It remains to be defined whether the transiently altered Treg cell/Tconv ratio has an effect on the peripheral homeostasis of these cells, especially when the cause of this atrophy is more protracted or more frequently taking place. Since thymus atrophy has been reported to occur in the context of chronic infections [8] and cancer [51, 52], it would be interesting to investigate the long-lasting consequences of thymus atrophy on Treg cell homeostasis and function in those settings.

## MATERIAL AND METHODS

### Animals

Foxp3^hCD2^ x Rag1^GFP^ reporter mice (C57BL/6 background) [30, 53] were bred and maintained at the animal facility of the Helmholtz Centre for Infection Research (Braunschweig, Germany). All mice were kept under specific pathogen-free conditions. In all experiments, female mice at the age of six weeks were used. All animal experiments were conducted in compliance with the German animal protection law (TierSchG BGBl. I S. 1105; 25.05.1998). Animals were housed and handled in accordance with good animal practice as defined by FELASA and the national animal welfare body GV-SOLAS. All animal experiments were approved by the Lower Saxony Committee on the Ethics of Animal Experiments as well as the Lower Saxony State Office of Consumer Protection and Food Safety under the permit number 33.19-42502-04-15/2058. All experiments were performed in accordance with the institutional, state, and federal guidelines.

### Virus

For the infection experiments, the mouse-adapted strain PR8M of IAV H1N1 was used. The virus was grown on MDCK cell line, harvested as a supernatant of the cultures, aliquoted and stored at −80°C. 666 ffu were given intranasally in 20 μl PBS to Foxp3^hCD2^ x Rag1^GFP^ reporter mice. Control mice received an intranasal application of 20 μl PBS.

### Cell sorting

CD4SP thymocyte subsets were sorted from thymi of control and infected mice. Thymic single-cell suspensions were enriched for CD4SP thymocytes by means of staining, using CD8α-APC antibody (53-6.7, purchased from Biolegend), followed by labeling with anti-APC microbeads (Miltenyi Biotec). DP and CD8SP thymocytes were depleted by applying the “depleteS” program of AutoMACS Pro (Miltenyi Biotec). Enriched CD4SP thymocytes were then stained with CD4-Pacific blue (RM4-5) fluorochrome-conjugated antibodies (BioLegend). FACS Aria IIu and FACS Aria II SORP (BD Biosciences) were employed for cell sorting of CD4SP thymocyte subsets.

### Antibodies and flow cytometry

Flow cytometric analysis was performed as described recently [54]. Cell suspensions isolated from thymi were labeled directly with the following fluorochrome-conjugated anti-mouse antibodies purchased from either BioLegend, BD Biosciences, or eBioscience: CD4-BV605 (RM4-5), CD8α-APC (53-6.7), CD25-PerCP-Cy5.5 (PC61.5), and human CD2-BV421 (RPA-2.10). Detection of active caspase 3/7 was accomplished using CellEvent™ Caspase-3/7 Red Detection Reagent (ThermoFisher Scientific). Sorted CD4SP Rag1^GFP+^ thymocytes were stained with CD5-PerCP-Cy5.5 (53-7.3) and Nur77-PE (12.14) antibody (both from eBiosciences). Exclusion of dead cells was done by using the LIVE/DEAD™ Fixable Blue stain kit (Invitrogen). Cells were acquired on LSR Fortessa™ flow cytometer (BD Biosciences), and data were analyzed with FlowJo software (BD Biosciences).

### Measurements of cytokines in the thymus

Thymi isolated from Foxp3^hCD2^ x Rag1^GFP^ reporter mice were homogenized and lysed using Bio-Plex Cell Lysis Kit (BioRad Laboratories), and the protein content of the thymic lysates was measured using Pierce™ BCA Protein Assay Kit (Thermo Scientific). Afterwards, equal protein amounts of the lysates (25 μg) were applied to the Bio-Plex Mouse Cytokine 23-plex Assay Kit (Bio-Rad Laboratories) and measured with the BioPlex200 system and BioPlex-Manager 6.2 according to manufacturer’s instructions. For the measurement of TGF-β levels in thymus, the same lysates were processed using LEGENDplex™ Mouse/Rat Free Active/Total TGF-β1 Assay (BioLegend) according to manufacturer’s instructions. Samples were then acquired using LSRFortessa™ flow cytometer, and data were analyzed with FlowJo software. TGF-β concentrations were normalized to thymic protein concentrations.

### Design of the mathematical model

The development of all major thymocyte populations is modeled as a set of Ordinary Differential Equations (ODEs). Three model structures were considered with different Treg cell differentiation pathways (see Supporting Information Fig. 4). The equations and parameters from the DP thymocyte compartment were directly taken from Thomas-Vaslin et al. [17]. The early proliferating DP (*eDP*) thymocytes were separated into generations (*G*_0_…*G*_Nmax_), while late DP (*lDP*) thymocytes are a unique population of cells dying or differentiating into SP thymocytes. The total DP thymocyte pool size was resized to 61.2 million cells as measured in the present study. The equations were adapted using a continuous number of divisions and a fraction of cells performing the last division, such that the proliferation rate can be made dependent on thymus size. By defining a maximum *N*_max_ of generations to be simulated, the equations for the eDP thymocyte pool follow:

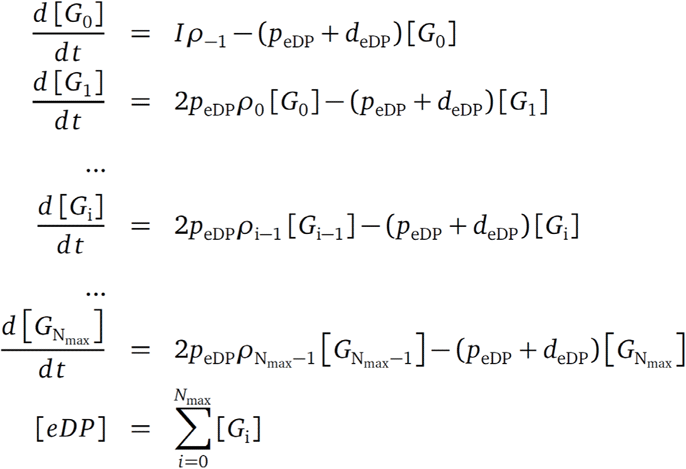

Where

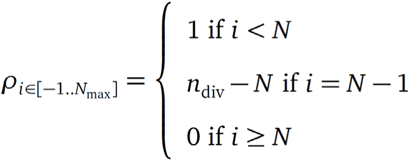

*I* denotes the inflow of cells from DN thymocytes. The population *G*_i_ represents eDP thymocytes that are proliferating, but did not complete yet the *i*+1^th^ cell cycle. With *N* = floor(*n*_div_) < N_max_, an average of *n*_div_ divisions is modeled with a fraction *n*_div_ - *N* of cells actually entering the *N*^th^ cycle (G_N_) while the remaining fraction exits to the lDP thymocyte stage. It appears as the fraction *ρ*_i_ of cells leaving *G*_i_ that are allowed to enter a new cell cycle, which is 1 for each division until *N*-2, a fraction for the *G*_N-1_, and 0 afterwards until *G*_Nmax_. The proliferation rate *p*_eDP_ denotes the rate of cells that succeed to reach the next generation, which appear as twice as many in the next generation. The flow of cells leaving the eDP pool to enter the lDP population is therefore:

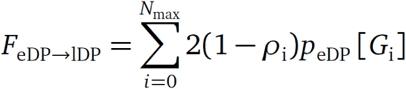

The differentiation between populations is taken as a linear process. The differentiation rates from lDP thymocytes into Tconv and CD8SP thymocytes are called *f*_2_ and *f*_5_, respectively. Depending on the model structure, Foxp3^+^CD25^-^ and Foxp3^-^CD25^+^ precursors arise from lDP or CD4SP thymocytes (Tconv) with rates *f*_3_ and *f*_4_ Therefore, the lDP thymocyte pool follows the equation:

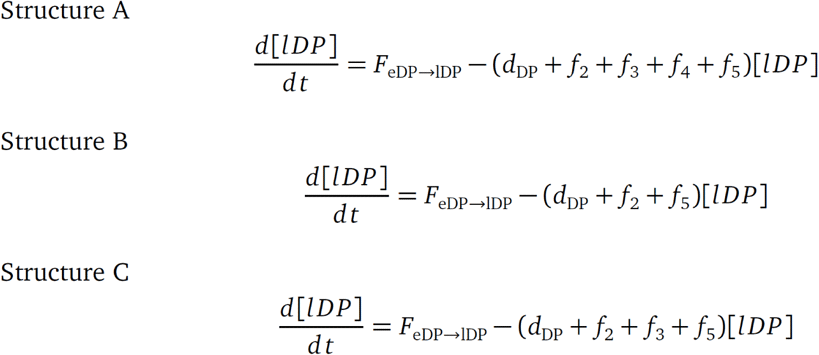

Although the eDP-lDP model is able to recover from atrophy quite fast, we realized that a compensatory mechanism on proliferation actually gives a slightly faster recovery and better fit, especially for hypothesis 1. This compensation was made as a ‘logistic growth’ term by using a dynamical number of divisions *N*_div_(t). We name *N*_div,init_ the initial number of divisions of eDP thymocytes at steady state and *K* a ‘carrying capacity’ of the thymus. T(t) represents the total number of T cells in the simulation, and α is scaled such that *N*_div_ = *N*_div,init_ at steady-state.

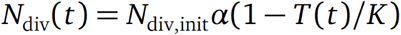

Death and output are the major driving mechanisms of SP thymocyte populations, which is compatible with an exponentially distributed residence time resulting from a single ODE. Proliferation, death, and thymic output are taken as constant rates named *p*_x_, *d*_x_, and *o*_x_, where x is the population name for SP thymocyte populations. *f*_6_ and *f*_7_ represent a constant differentiation rate between Treg cell precursors to mature Treg cells.

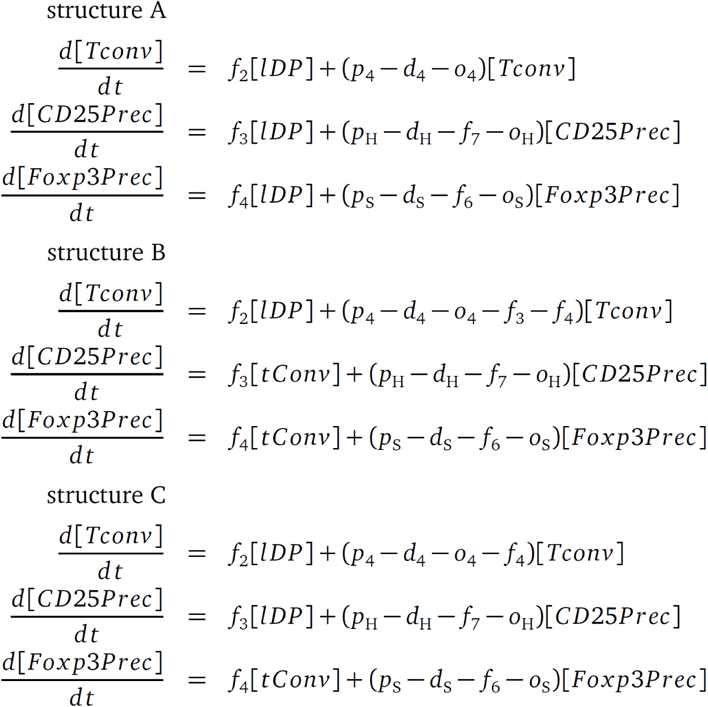

And for all model structures,

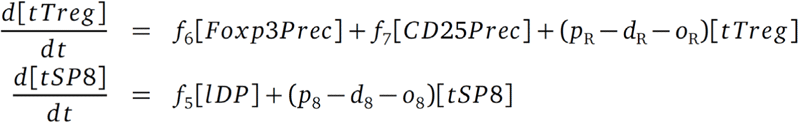

#### Parameter values

The parameter values for Tconv and CD8SP thymocyte dynamics were determined according to two constraints: (i) The proliferation rate is derived from the death and output rates to maintain experimental steady-state values, and (ii) the residence time of cells in the CD4SP and CD8SP thymocyte compartment was estimated to be 96 hours, which is calibrated as 1/(*o*_x_+*d*_x_-*p*_x_) in the model [55]. This suffices to determine the differentiation rate of lDP into CD4SP and CD8SP thymocytes, and to estimate the proliferation rate as a function of the death and output parameters. The remaining parameters (death and output of Tconv or CD8SP thymocytes, respectively) were completely insensitive, and the curves during atrophy were independent of their values (data not shown). Therefore, death and output rates were fixed with realistic values.

For the Treg cell precursors and mature Treg cell populations, there is no clear consensus on their residence time in the thymus, so only the steady state constraint was applied and the proliferation rate p_x_ of each population is taken as a function of its death and output rates *d*_x_ and *o*_x_, which are also insensitive. The death rates *d*_x_ were estimated following two experimental constraints: (i) CD25+Foxp3^-^ precursors die on average 3.67 times more than CD25^-^Foxp3^+^ precursors, as measured by Annexin V staining in Owen et al. [6], and (ii) mature tTreg cells die on average 1.9 times more than Tconv [56]. We assumed that CD25^-^Foxp3^+^ Treg cell precursors and mature Treg cells die with a similar strength.

Four main parameters are unknown: i) conversion rate from lDP thymocytes to CD25^+^Foxp3^-^ precursors, ii) conversion rate from lDP thymocytes to CD25^-^Foxp3^+^ precursors, iii) conversion rate from CD25^+^Foxp3^-^ precursors to mature tTreg cells and iv) conversion rate from CD25^-^Foxp3^+^ precursors to mature tTreg cells. Similarly as for Tconv and CD8SP thymocytes, the death and output parameters were insensitive and could be fixed without impact. The list of parameters fixed in the system are summarized in Table 1.

**Table 1:**
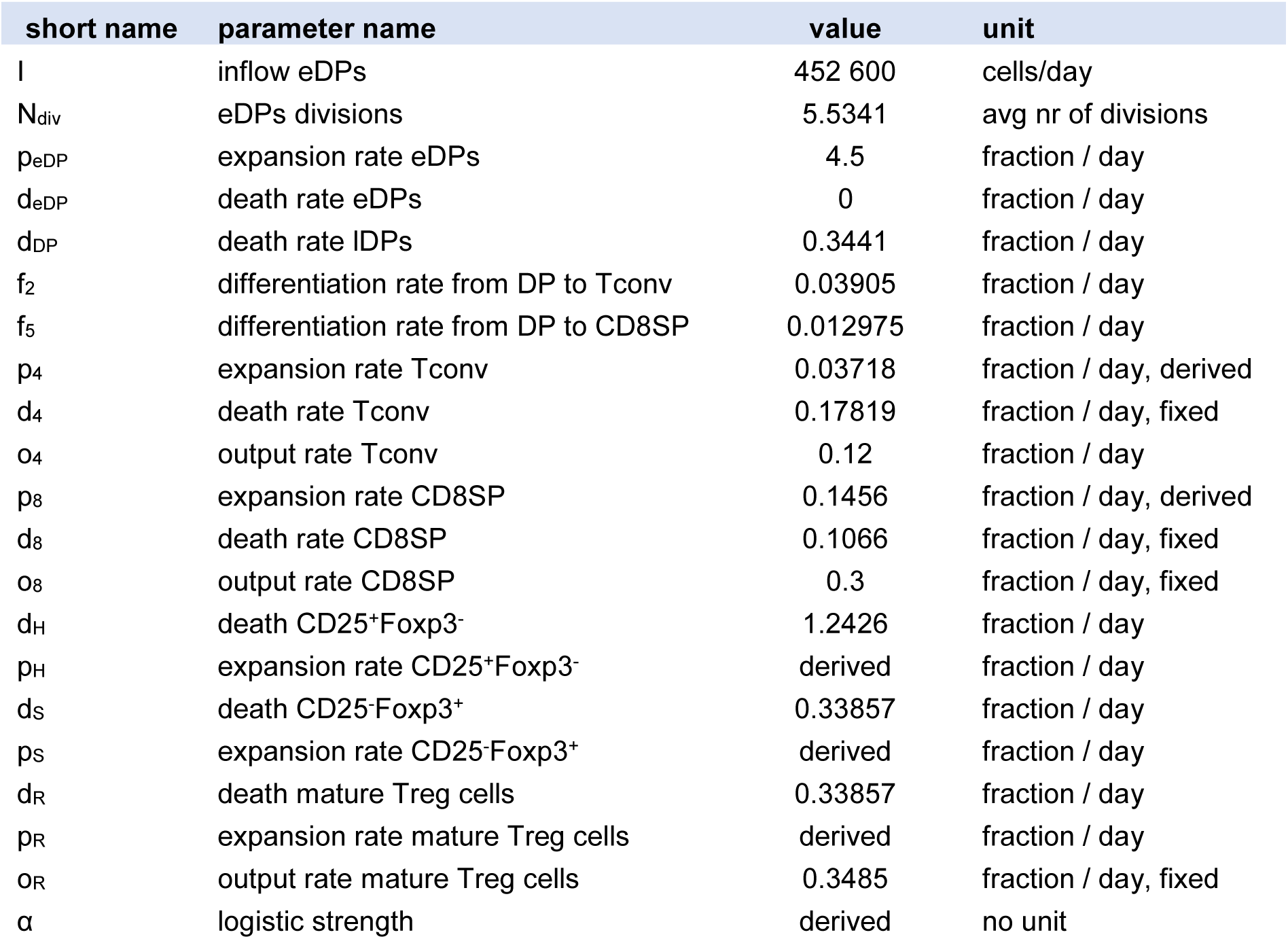
Parameter values fixed before optimization. Proliferation rates are derived from other parameter values under steady-state constraint. The proliferation of Treg cell precursors and Treg cells will depend on the differentiation rates that are unknown and included in the fittings, therefore called ‘derived’.

#### Parameter optimization

The compatibility of different atrophy models (model structure and atrophy hypotheses) was assessed by parameter optimization (‘fitting’) using Stochastic Ranking Evolution Strategy algorithm [57] and by comparing simulations with the experimental dataset. The used cost function is the Residual Sum of Squares (RSS) after normalizing each population by its average value along all time-points, such that each population has a balanced contribution to the total cost. Best fitted parameter values are shown in Table 2, together with the RSS.

**Table 2:**
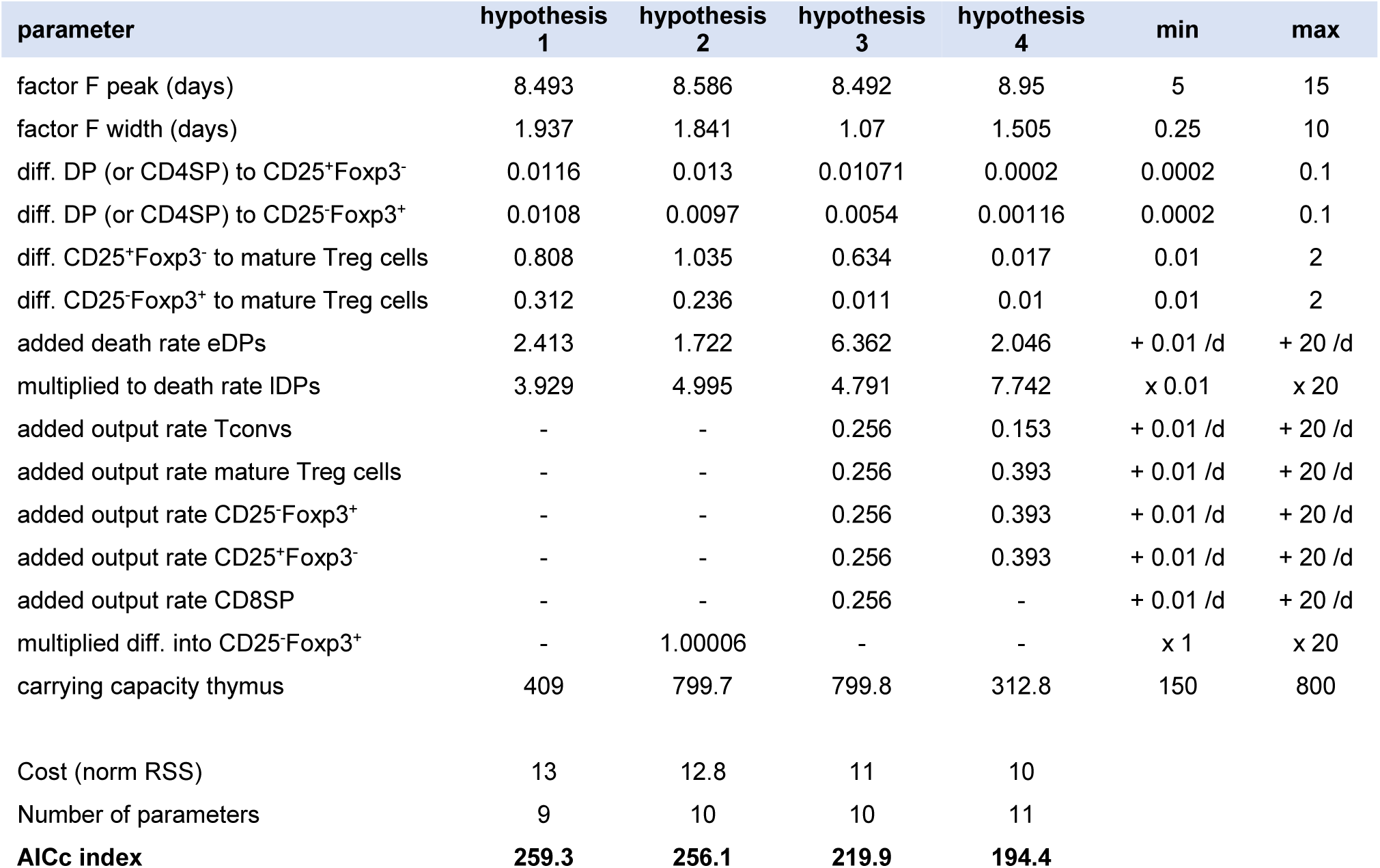
Parameter boundaries, best parameter sets and RSS costs. obtained during the fitting of each hypothesis for model structure B and corresponding to the curves shown in Figure 6.

The AICc compares the quality/cost of models with different number of parameters. A minimal AICc value defines the model with the best fit and a minimal number of parameters. A difference of 2 in the AICc value is taken as significant improvement as a rule of thumb [58]. We assume a Gaussian distribution of noise in the *n*=169 data points, but with different strength per data point. The AICc formula is derived from the likelihood *L* of the simulated points *x*_*θ,j*_ with the best parameter set θ as following, with *k* parameters,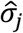and 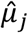 the estimated variance and mean of each data point *j*:

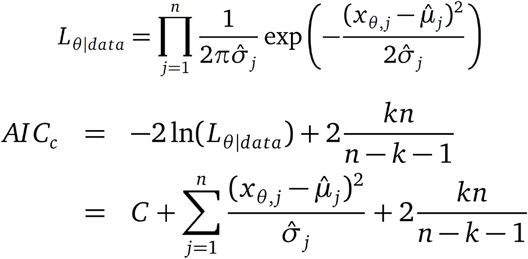

C represents a constant that is neglected when comparing the difference of AICc values between models, and is ignored in the calculation.

The experimental dataset, details of modeling hypotheses and parameter estimation results for all model structures are given in Supporting Information ‘Mathematical Model’. The simulation code, written in C++ using the Moonfit framework [59], was deposited at https://gitlab.com/Moonfit/Balthyse.

### TCR profiling

For the TCR repertoire analysis, Treg cells and Tconv were sorted from thymi of control or IAV-infected mice as Rag1^GFP+^CD25^+^Foxp3^hCD2+^, Rag1^GFP-^CD25^+^Foxp3^hCD2+^, Rag1^GFP+^CD25^-^Foxp3^hCD2-^, and Rag1^GFP-^CD25^-^Foxp3^hCD2-^ CD4SP thymocyte subsets. A 5’RACE cDNA synthesis, followed by a PCR employing a mTCRβ constant region reverse primer and a UPM (universal primer mix), was performed. In brief, RNA extraction from sorted cells was performed by using RNeasy Plus Micro kit (Qiagen). The cDNA was generated by employing ingredients from the TAKARA cDNA Synthesis Kit (TAKARA Bio Europe), based on the manufacturer’s protocol. A mixture from 1 μl 12 μM OligodT and 4.5 μl RNA was incubated for 3 min at 72°C, followed by 2 min at 42°C and placed on ice immediately. Directly, 4.5 μl RACE-Mastermix (1X First-Strand-buffer, 2.5 mM DTT, 1 mM dNTP mix, 0.6 μM SMARTer IIA Oligonucleotide, 10 U RNase Inhibitor and 50 U SMARTScribe Reverse Transcriptase) was added, and the PCR program was continued for 90 min at 42°C and 10 min at 72°C. Subsequently, the DNA-amplicons for Illumina sequencing were generated by using Advantage 2 PCR Kit (TAKARA Bio Europe), employing a mTCRβ reverse primer and a Universal primer mix (UPM). The fully generated cDNA (10 μl) was processed. After agarose-electrophoresis, the DNA was purified using the Qiaquick Gel Extraction Kit (Qiagen). The purified product of each sample was indexed by an indexing-PCR with Nextera-Primers. Finally, a pool of the samples was sequenced, employing Illumina MiSeq v2 Reagent Kit (Illumina). For the analysis of TCR sequencing data, samples were aligned to the international immunogenetics information system (IMGT) database from 22.05.2018, using MiXCR software [60]. Then, the TCR diversity analysis (inverse Simpson’s index) was performed by employing VDJTools [61]. The analyses were done to the complete data sets, and to thymus samples down-sampled to 20,000 counts (i.e. productive reads) after assessment of read-numbers in all samples. Sequencing data were deposited at NCBI GEO with accession number GSE139499.

### Statistical analysis

The GraphPad Prism software v7.0 was used to perform all statistical analyses. Data are presented as mean ± or + standard deviation (SD). For comparison of unmatched groups, the two-tailed Mann-Whitney test was performed, and the p-values were calculated with the long-rank test (Mantel-Cox). A p-value below 0.05 was considered as significant; * p<0.05; ** p<0.01; *** p<0.001; **** p<0.0001; ns (not significant).

## Supporting information

All Supplementary Information

## Abbreviations

AICc: corrected Akaike Information Criterion
DP: double-positive
dpi: days post infection
IAV: influenza A virus
lDP: late double-positive
ODE: ordinary differential equation
SP: single-positive
Tconv: conventional T cell
TCR: T cell receptor
Treg: regulatory T cell

## Acknowledgements

We thank Lothar Gröbe for cell sorting, Maria Ebel for technical support and Marcus Gereke. This work was supported by PhD scholarship programme no. 57129429 of the German Academic Exchange Service (DAAD) (to Y.E.). P.A.R. was supported by the Human Frontier Science Program (RGP0033/2015).

## Conflicts of interest

The authors declare no commercial or financial conflict of interest.

## Notes

https://gitlab.com/Moonfit/Balthyse

### Competing Interest Statement

The authors have declared no competing interest.

### Summary of Updates

Solved a problem in the AIC calculation, updated with new AICc values and recalculated Supp. Figure S8.

