## Supplementary material for "Influenza A virus-induced thymus atrophy differentially affects dynamics of conventional and regulatory T cell development": All Supplementary Information

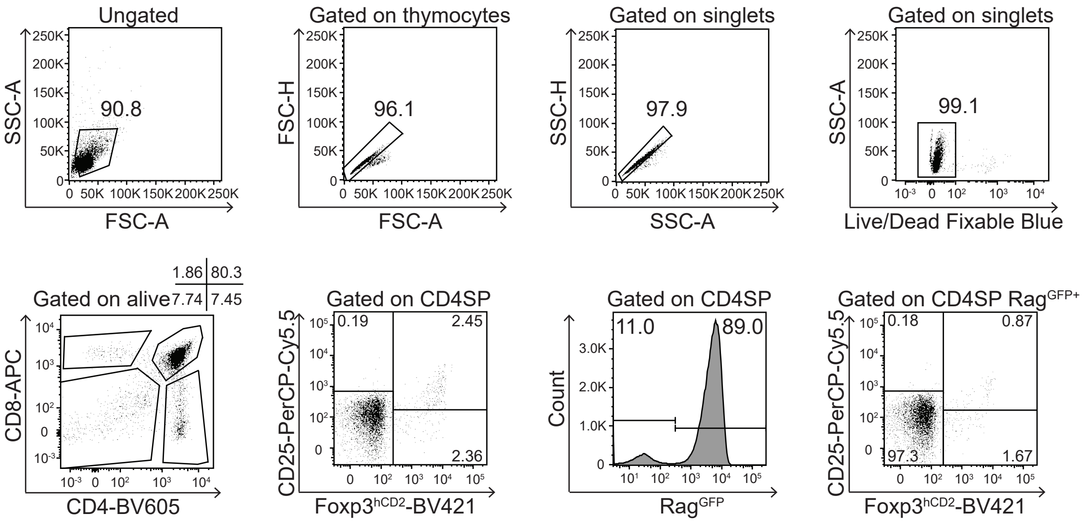

**Supporting Information Figure 1. Gating strategy for the identification of total and newly generated Treg cells and their precursors in the thymus.** Single-cell suspensions were prepared from thymi of PBS-treated Foxp3$^{hCD2}$xRag1$^{GFP}$ double reporter mice and analyzed by flow cytometry. Expression of CD4 and CD8 in thymocytes is shown after excluding doublets and dead cells. Treg cells and their precursors are displayed either gated on total CD4SP thymocytes or newly generated Rag1$^{GFP+}$ CD4SP thymocytes. Numbers indicate frequencies of cells in the indicated gates.

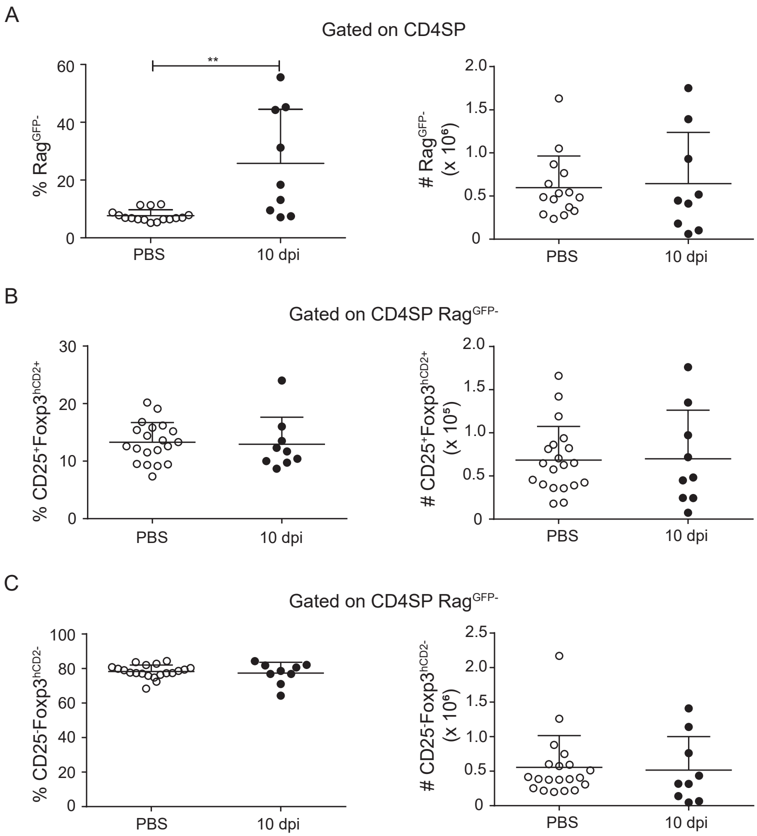

**Supporting Information Figure 2. Recirculating Treg cells or Tconv are unaffected during IAV infection.** Foxp3$^{hCD2}$xRag1$^{GFP}$ double reporter mice were infected with IAV and control mice received PBS. Thymi were analyzed at different dpi. Scatterplots summarize frequencies (left) and absolute numbers (right) of CD4SP Rag1$^{GFP-}$ thymocytes (A), Rag1$^{GFP-}$CD25$^{+}$Foxp3$^{+}$ Treg cells (B), and Rag1$^{GFP-}$CD25$^{-}$Foxp3$^{-}$ Tconv (C) from PBS-treated (open circles) and IAV-infected mice (filled circles) (10 dpi). Data were pooled from three independent experiments with 9-20 mice per group and presented as mean + SD. Each dot represents an individual mouse. Mann-Whitney test was used to test for statistical significance.

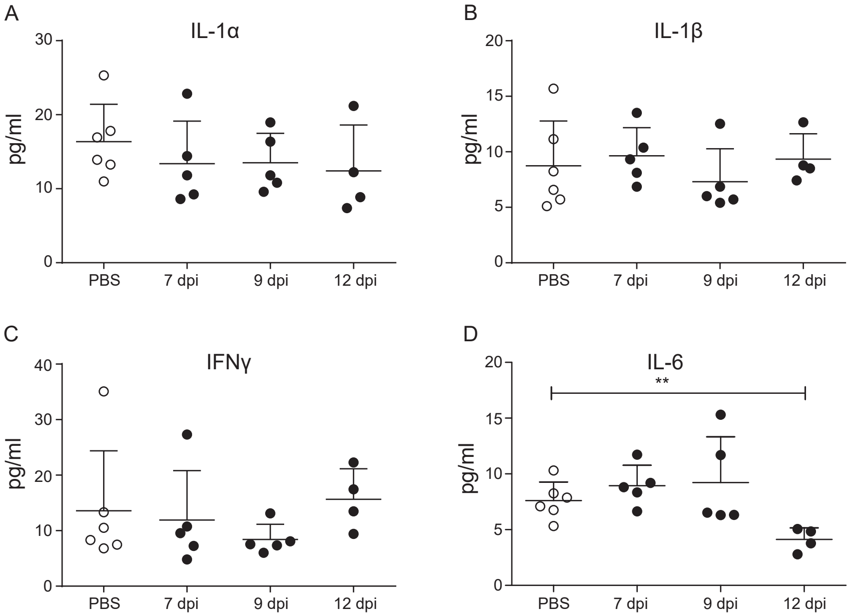

**Supporting Information Figure 3. Cytokine levels in thymi post IAV infection.** $Foxp3^{hCD2}xRag1^{GFP}$ double reporter mice were infected with IAV and control mice received PBS. Thymus lysates were analyzed at different dpi. Concentrations of IL-1α, IL-1β, IFNγ, and IL-6 were measured by Bio-plex 23-plex mouse kit. Scatterplots summarize IL-1α (A), IL-1β (B), IFNγ (C), and IL-6 (D) concentration in thymi from PBS-treated (open circles) and IAV-infected mice (filled circles) at indicated dpi. Data were pooled from two independent experiments with 4-6 mice per group and presented as mean + SD. Each dot represents an individual mouse. Mann-Whitney test was used to test for statistical significance.

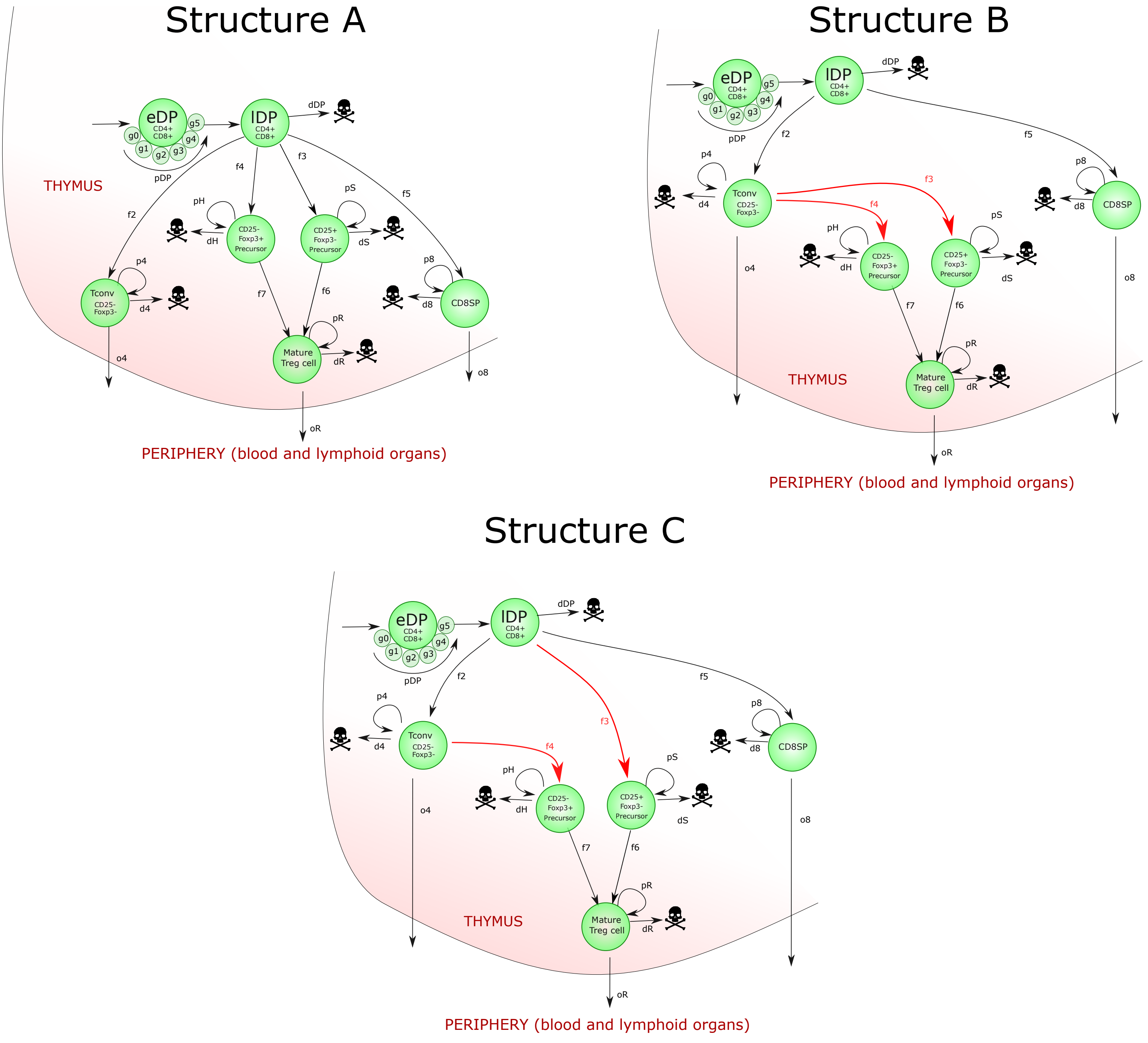

**Supporting Information Figure 4. Mathematical models considered for thymic Treg development.** Treg cell precursors either arise directly from DP thymocytes (structure A), or first become CD4SP thymocytes (structure B). Alternately, only CD25$^-$Foxp3$^+$ Treg cell precursors arise from CD4SP thymocytes (structure C).

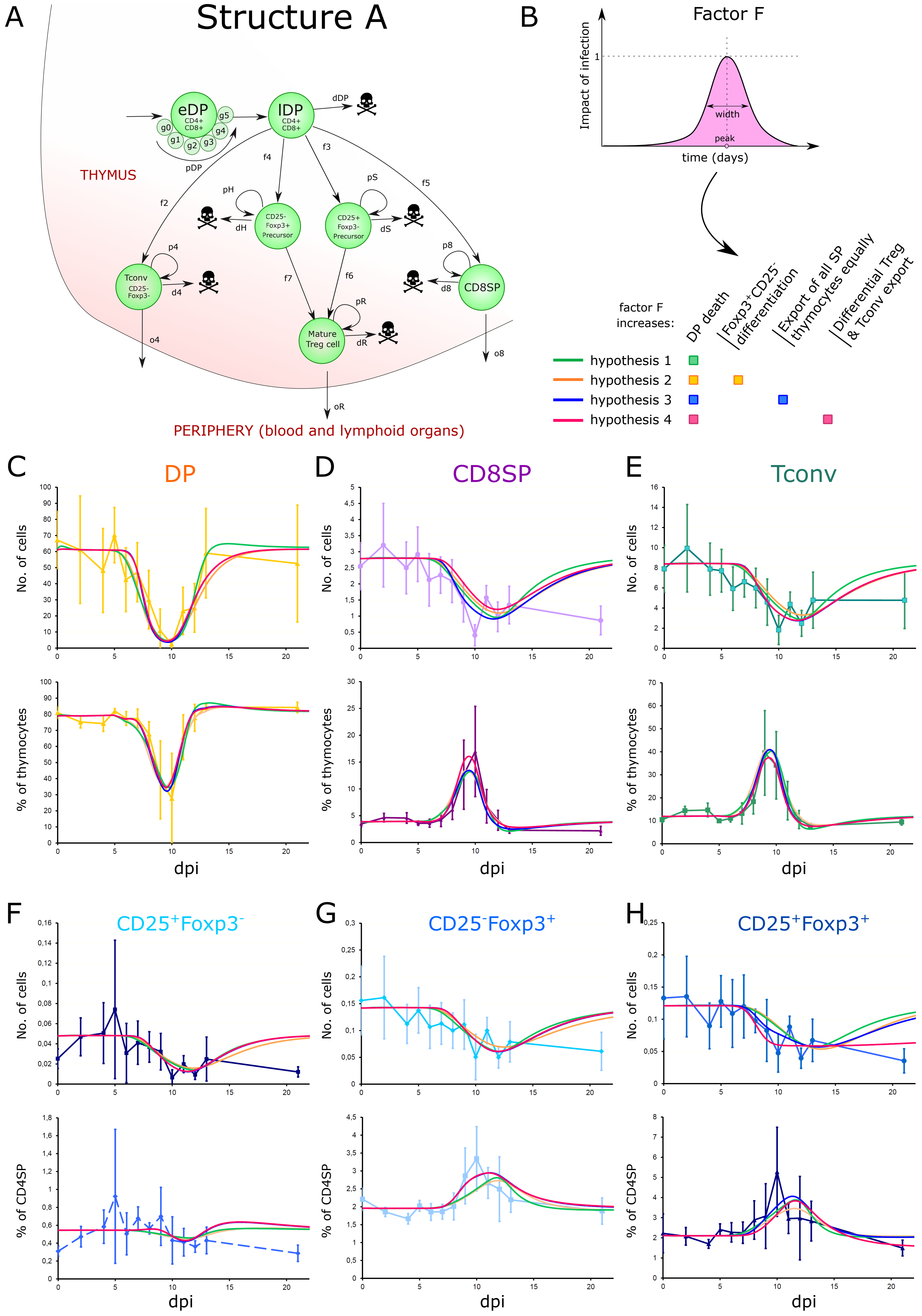

**Supporting Information Figure 5. Curves of the best fits for model structure A under different model hypotheses of IAV-induced thymus atrophy.** (A) Model structure A. (B) Tested hypotheses. (C-H) Dynamics of thymic atrophy under the hypothesis of increased DP thymocyte death (hypothesis 1), together with increased *de novo* generation of $CD25^{-}Foxp3^{+}$ precursors (hypothesis 2), together with equally increased export of all SP thymocytes (hypothesis 3), or together with differentially increased export of Tconv or Treg cells (hypothesis 4). Best curves shown as absolute numbers (top) or percent (bottom).

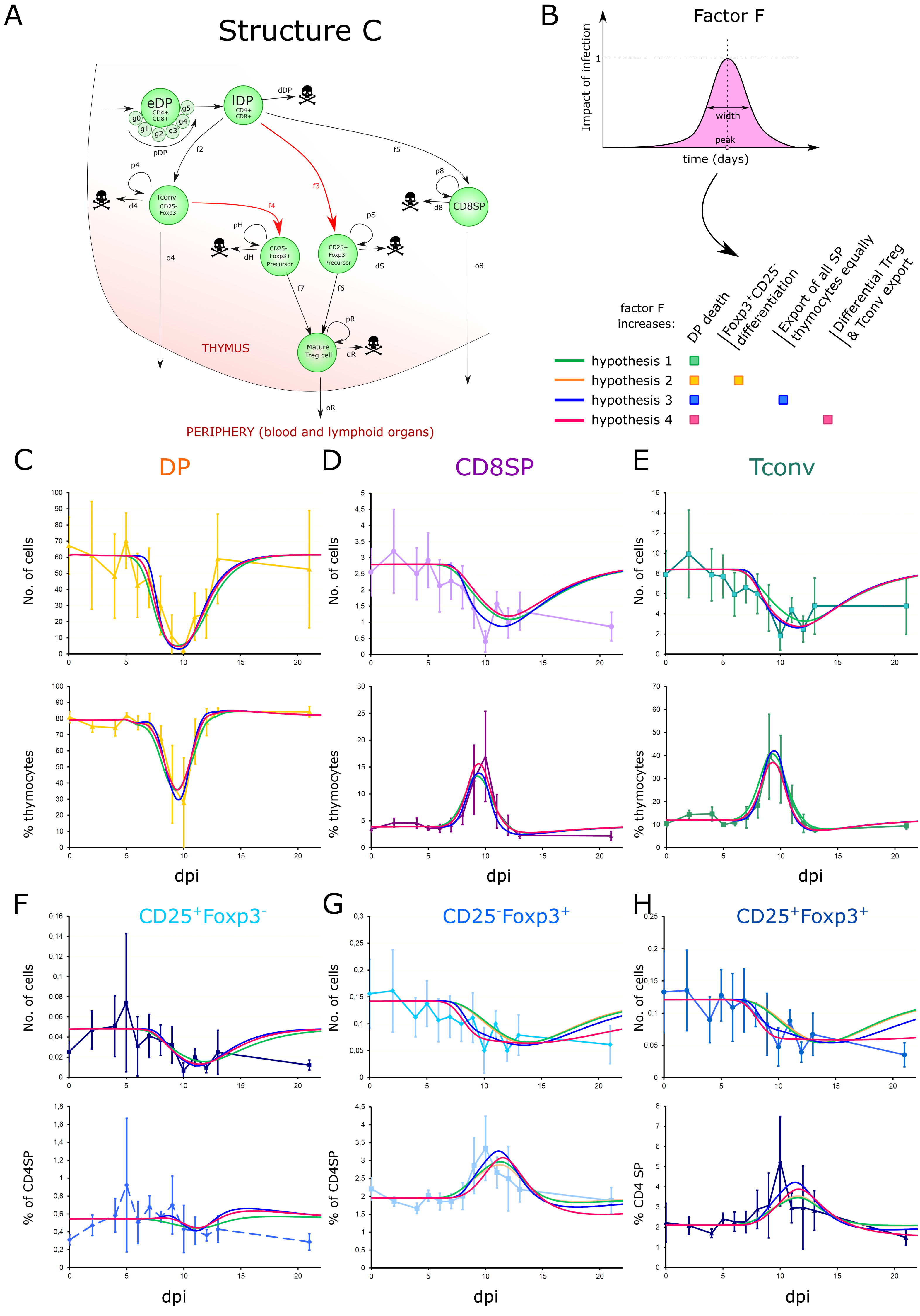

**Supporting Information Figure 6. Curves of the best fits for model structure C under different model hypotheses of IAV-induced thymus atrophy.** (A) Model structure C. (B) Tested hypotheses. (C-H) Dynamics of thymic atrophy under the hypothesis of increased DP thymocyte death (hypothesis 1), together with increased *de novo* generation of CD25$^-$Foxp3$^+$ precursors (hypothesis 2), together with equally increased export of all SP thymocytes (hypothesis 3), or together with differentially increased export of Tconv or Treg cells (hypothesis 4). Best curves shown as absolute numbers (top) or percent (bottom).

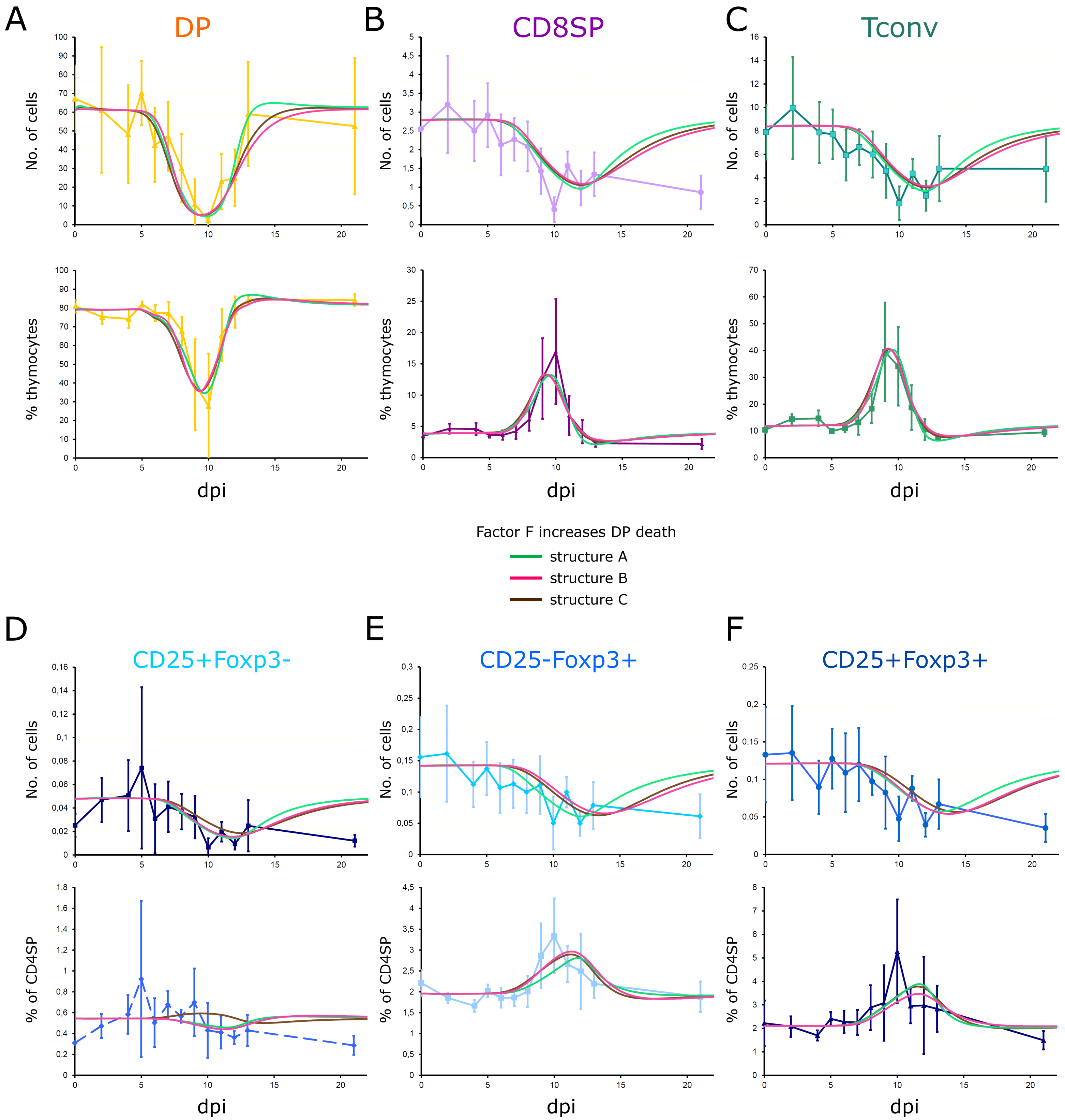

**Supporting Information Figure 7. Shrinkage of DP thymocyte population is the dominant factor on SP thymocyte population dynamics during IAV-induced thymus atrophy.** Dynamics of thymus atrophy for each model structure according to their best fit under increased eDP and lDP thymocyte death (hypothesis 1 in Figure 6). Dynamics of the main thymocyte populations are shown in each panel as absolute numbers (top) or percent (bottom).

| Controls without DP death: | Model structure A | | Model structure B | | Model structure C | |
|---|---|---|---|---|---|---|
| Factor F **increases** | | | | | | |
| De novo CD25-Foxp3+ Treg cell prec. generation | X | | X | | X | |
| Export of all SP cells equally | | X | | X | | X |
| | | | **Quality of the best fit** | | | |
| **Cost best set (normalized RSS)** | **43.9** | **45.3** | **42.4** | **33.5** | **45.9** | **41.5** |
| Number of unknown parameters | **8** | **8** | **8** | **8** | **10** | **10** |
| AICc index | **937.6** | **914.8** | **858.4** | **664.8** | **1065** | **903.2** |
| Comparison (Supporting Information Figure 9) | | | | | | |
| Cost **with** DP death | **12.4** | **11.5** | **12.8** | **11.0** | **12.5** | **11.2** |
| AICc index **with** DP death | **247.3** | **235.3** | **256.1** | **219.9** | **245.9** | **230.2** |

**Supporting Information Figure 8. Cost (normalized RSS) and AICc index for each model structure under hypotheses where factor F does not impact on DP thymocyte death.**

| | Model structure A | | | | Model structure B | | | | Model structure C | | | | Min | Max |
|---|---|---|---|---|---|---|---|---|---|---|---|---|---|---|
| Factor F **increases** | hyp 1 | hyp 2 | hyp 3 | hyp 4 | hyp 1 | hyp 2 | hyp 3 | hyp 4 | hyp 1 | hyp 2 | hyp 3 | hyp 4 | | |
| DP death | X | X | X | X | X | X | X | X | X | X | X | X | | |
| De novo CD25$^-$Foxp3$^+$ Treg cell prec. generation | | X | | | | X | | | | X | | | | |
| Export of all SP thymocytes equally | | | X | | | | X | | | | X | | | |
| Export of Tconv and Treg cells differentially | | | | X | | | | X | | | | X | | |
| **Quality of the best fit** | | | | | | | | | | | | | | |
| Cost of best parameter set (norm RSS) | **12.3** | **12.4** | **11.5** | **11.2** | **13.0** | **12.8** | **11.0** | **10.0** | **12.5** | **12.5** | **11.2** | **10.9** | | |
| Number of unknown parameters | **9** | **10** | **10** | **11** | **9** | **10** | **10** | **11** | **9** | **10** | **10** | **11** | | |
| AICc index | **251.0** | **247.3** | **235.3** | **236.1** | **259.3** | **256.1** | **219.9** | **194.4** | **244.3** | **245.9** | **230.2** | **233.1** | | |
| **Fitted parameter values** | | | | | | | | | | | | | **Min** | **Max** |
| factor F peak (days) | 9.211 | 8.504 | 8.333 | 8.389 | 8.493 | 8.586 | 8.492 | 8.950 | 8.727 | 8.695 | 8.439 | 8.471 | 5 | 15 |
| factor F width (days) | 1.857 | 1.723 | 1.123 | 1.232 | 1.937 | 1.841 | 1.070 | 1.505 | 1.710 | 1.709 | 0.944 | 1.377 | 0.250 | 10.000 |
| diff. rate from DP (or CD4SP) into CD25$^+$Foxp3$^-$ precursors | 0.000263 | 0.000282 | 0.000414 | 0.000423 | 0.0116 | 0.0130 | 0.01071 | 0.000200 | 0.000286 | 0.000287 | 0.000465 | 0.000401 | 0.0002 | 0.1 |
| diff. rate from DP (or CD4SP) into CD25$^-$Foxp3$^+$ precursors | 0.000504 | 0.000473 | 0.000657 | 0.000669 | 0.0108 | 0.0097 | 0.00540 | 0.001160 | 0.00923 | 0.01071 | 0.00534 | 0.00171 | 0.0002 | 0.1 |
| diff. rate from CD25$^+$Foxp3$^-$ precursors to mature Treg cells | 0.258 | 1.103 | 0.544 | 0.0106 | 0.808 | 1.035 | 0.634 | 0.017 | 1.107 | 0.864 | 0.069 | 0.012 | 0.010 | 2.000 |
| diff. rate from CD25$^-$Foxp3$^+$ precursors to mature Treg cells | 0.498 | 0.010 | 0.010 | 0.010 | 0.312 | 0.236 | 0.011 | 0.010 | 0.208 | 0.294 | 0.553 | 0.039 | 0.010 | 2.000 |
| additive death rate of early DP thymocytes | 2.459 | 2.127 | 8.700 | 4.799 | 2.413 | 1.722 | 6.362 | 2.046 | 1.504 | 1.604 | 11.158 | 3.140 | + 0.01 /d | + 20 /d |
| multiplicative death rate of late DP thymocytes | 16.83 | 4.35 | 3.816 | 3.800 | 3.929 | 4.995 | 4.791 | 7.742 | 6.181 | 5.864 | 4.780 | 4.109 | x 0.01 | x 20 |
| additive output rate of Tconv cells | - | - | 0.175 | 0.167 | - | - | 0.256 | 0.153 | - | - | 0.268 | 0.148 | + 0.01 /d | + 20 /d |
| additive output rate of mature Treg cells | - | - | 0.175 | 0.332 | - | - | 0.256 | 0.393 | - | - | 0.268 | 0.288 | + 0.01 /d | + 20 /d |
| additive output rate of CD25$^-$Foxp3$^+$ precursors | - | - | 0.175 | 0.332 | - | - | 0.256 | 0.393 | - | - | 0.268 | 0.288 | + 0.01 /d | + 20 /d |
| additive output rate of CD25$^+$Foxp3$^-$ precursors | - | - | 0.175 | 0.332 | - | - | 0.256 | 0.393 | - | - | 0.268 | 0.288 | + 0.01 /d | + 20 /d |
| additive output rate of CD8SP thymocytes | - | - | 0.175 | - | - | - | 0.256 | - | - | - | 0.268 | - | + 0.01 /d | + 20 /d |
| multiplicative diff. rate from DP (or CD4SP) to CD25$^-$Foxp3$^+$ prec. | - | 1.00007 | - | - | - | 1.00006 | - | - | - | 1 | - | - | x 1 | x 20 |
| carrying capacity of the thymus (*) | 162.8 | 800.0 | 797.9 | 796.9 | 409.0 | 799.7 | 799.8 | 312.8 | 799.0 | 799.8 | 799.8 | 797.8 | 150 | 800 |
| **Derived parameter values (steady state)** | | | | | | | | | | | | | | |
| prolif. rate of CD25$^+$Foxp3$^-$ precursors | 1.194 | 2.016 | 1.304 | 0.760 | 0.0167 | 0.0002306 | 0.0018 | 1.224 | 2.015 | 1.771 | 0.768 | 0.786 | | |
| prolif. rate of CD25$^-$Foxp3$^+$ precursors | 0.638 | 0.162 | 0.090 | 0.085 | 0.0142 | 0.0000501 | 0.030 | 0.280 | 0.001017 | 0.000035 | 0.576 | 0.276 | | |
| prolif. rate of mature Treg cells | 0.000272 | 0.2386 | 0.460 | 0.671 | 0.000836 | 0 | 0.423 | 0.669 | 0.00496 | 0.000151 | 0.01040 | 0.637 | | |
| strength of logistic control of DP proliferation | 1.904 | 1.107 | 1.107 | 1.107 | 1.233 | 1.107 | 1.107 | 1.328 | 1.107 | 1.107 | 1.107 | 1.107 | | |
| **Predicted half-life of Treg cell and Treg cell precursor populations (qualitative validation)** | | | | | | | | | | | | | | |
| CD25$^+$Foxp3$^-$ precursors half-life | 3.3 | | 2.1 | 2.0 | | | **0.5** | **28.6** | | | 1.8 | 2.1 | | |
| CD25$^-$Foxp3$^+$ precursors half-life | 5.0 | | 3.9 | 3.8 | | | **3.1** | **14.6** | | | 3.2 | **9.9** | | |
| mature Treg cells half-life | 1.5 | | 4.4 | **62.5** | | | **3.8** | **54.2** | | | 1.5 | **19.9** | | |

**Supporting Information Figure 9. Best parameter values and cost for each model structure and hypothesis.** The cost (RSS) and AICc index are shown, together with the best fitted parameter values and the derived parameters to ensure steady state (proliferation). Further, the predicted half-life of Treg cell and Treg cell precursor populations is shown for the best sets as a qualitative validation. Red numbers are those with inconsistent biological meaning. The green column shows the best parameter set that satisfies physiological half-lives, according to the cost and AICc index.

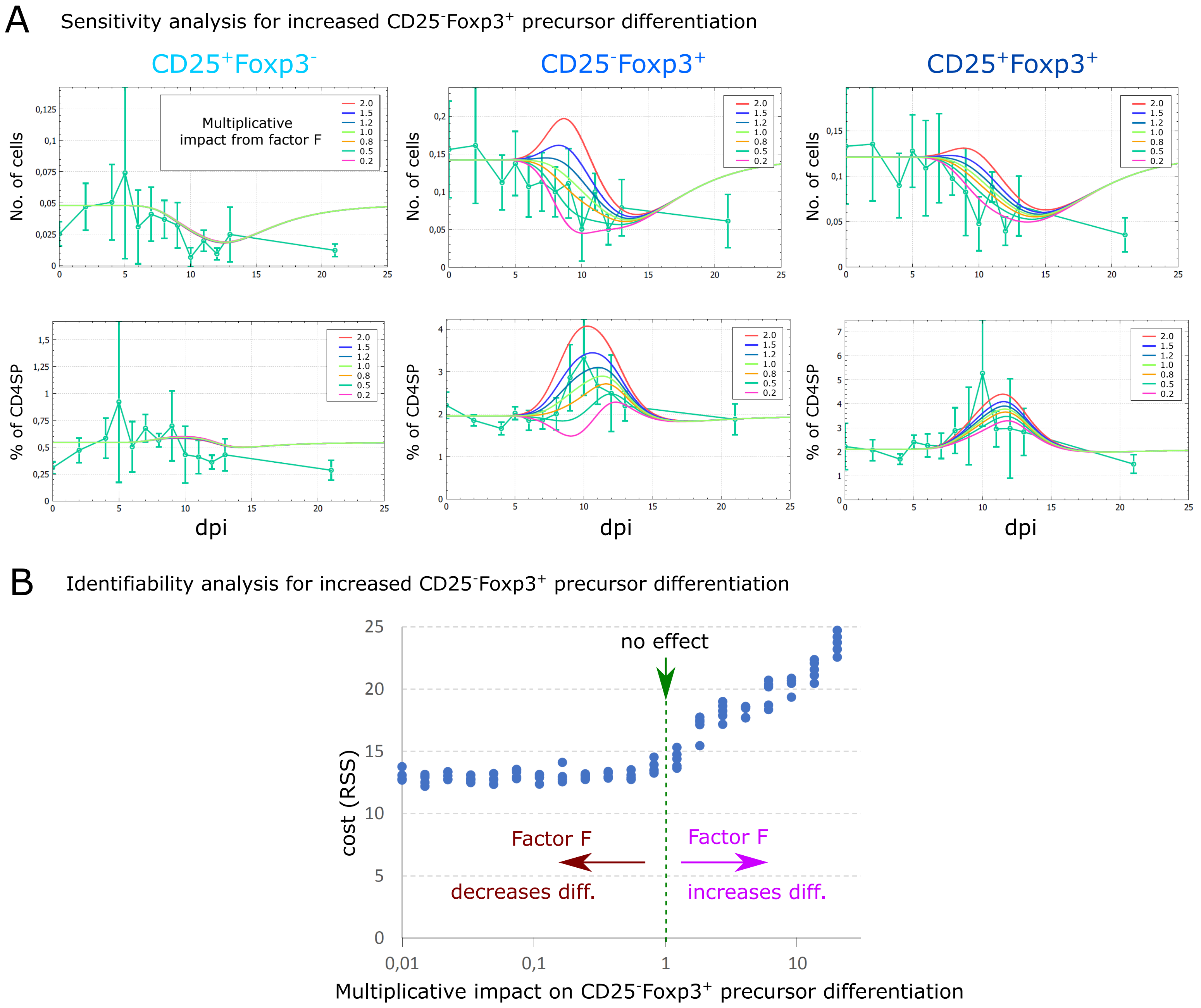

**Supporting Information Figure 10. Impact of *de novo* generation of CD25⁻Foxp3⁺ Treg cell precursors on the dynamics of Treg cell and Treg cell precursor populations during IAV-induced thymus atrophy.** (A) From the best parameter set of thymus atrophy with increased DP thymocyte death (hypothesis 1), impact of factor F increasing or decreasing *de novo* generation of $CD25^{-}Foxp3^{+}$ precursors. (B) Identifiability analysis: for each possible value of increased or decreased *de novo* $CD25^{-}Foxp3^{+}$ precursor generation, the cost of parameter estimation is given when factor F modulated DP thymocyte death and *de novo* generation of $CD25^{-}Foxp3^{+}$ precursors with this strength. Factor F increasing *de novo* generation of $CD25^{-}Foxp3^{+}$ precursors leads to a worsened cost while a decreased *de novo* generation of $CD25^{-}Foxp3^{+}$ precursors is not significantly improving the cost.

### Supporting Information 'Mathematical Model'

The datasets used to design and calibrate the model are presented in the following sections.

**Steady-state size of major thymocyte populations:** The steady-state size of each major thymocyte population was determined in 6 to 7 weeks old age-matched Foxp3$^{hCD2}$ x Rag1$^{GFP}$ reporter mice (C57BL/6 background).

| Population | Abs Number | % of T cells | Rag+ | Rag- | %Rag- |
|---|---|---|---|---|---|
| DN1 + $lin^+$ (1) | | 0,771 % ±0,251 | 553 300 (2) | | |
| DN2 | | 0,133 % ±0,061 | 97 390 (2) | | |
| DN3 | | 1,61 % ±0,541 | 1 217 000 (2) | | |
| DN4 | | 2,60 % ±1,20 | 1 929 000 (2) | | |
| DP total | | 78,2 % ±4,85 | 61 240 000 | | |
| CD8SP | 3 016 000 | 4,11 % ±0,87 | 2 787 000 | 229 000 | 7,54 ±3,2 |
| Tconv (3) | 8 816 000 | 12,0 % ±2,79 | 8 387 000 | 429 200 | 4,87 ±0,82 |
| $CD25^+Foxp3^-$ | 50 270 | 0,069 % ±0,042 | 47 930 | 2 350 | 6,22 ±4,1 |
| $CD25^-Foxp3^+$ | 180 900 | 0,239 % ±0,045 | 142 000 | 38 960 | 21,0 ±5,6 |
| Mature Treg cells | 195 000 | 0,257 % ±0,070 | 121 100 | 73 890 | 36,5 ±7,5 |
| Total T cells | 77 300 000 | 100 % | | | |
| Total thymus cells | 77 560 000 ± 29 400 000 | | | | |

**Table 'Steady-state size of major thymocyte populations'** Calculated as the average of the first four time-points of the kinetics (before impact of infection, days 0, 2, 4 and 5; n=21 mice). Rag1$^{GFP+}$ newly generated and Rag1$^{GFP-}$ resident or recirculating cells were discriminated. **(1)** It has to be noted that the population of DN1 thymocytes is composed of precursors of DN2 thymocytes (the 'real' DN1), but also a majority of cells that can express TCRβ [1], CD19 (B cells) or NK cell markers [2,3]. Therefore, without additional markers (lin), DN1 thymocytes cannot be unequivocally defined as committed T cells precursors. Rescaled for a total thymocyte size of 77 M cells, Tan et al. [1] estimated the number of 'real' DN1 thymocytes as only 20,000, while Porritt et al. [2] estimated 250,000 cells for a thymus of 250 M cells, that would lead to around 77,000 cells here. In the simulations, DN thymocytes are constant and we only consider a constant inflow of cells into eDP thymocytes. **(2)** DN thymocytes are mixed between Rag1$^{GFP+}$ and Rag1$^{GFP-}$ cells, with an increasing frequency of Rag1$^{GFP+}$ cells along DN1 to DN4 thymocyte stages, reaching 100 % at the DP thymocyte stage. The number of Rag1$^{GFP+}$ and Rag1$^{GFP-}$ cells is only shown for later thymocyte stages because Rag1$^{GFP+}$ has been expressed before in those cells and can be used as a timer for longer-term residency or recirculation. **(3)** Only CD25$^-$Foxp3$^-$ CD4SP thymocytes are counted as 'Tconv'. The CD4 total would be the sum of Tconv, mature Treg cells and Treg cell precursors.

**Dynamics of major thymocyte populations upon IAV infection:** The dynamics of the main thymocyte populations were determined at several time points after IAV infection. Although the thymus is involuting with age, in the present study age-dependent thymus atrophy was neglected as this involution is negligible at the scale of the experiments presented here. Further, the shrinkage as absolute numbers is expected to be around 10-15 % over one month at that age [4], and the effects of the IAV infection are concentrated on 3 to 5 days, between 8 to 12 dpi. However, the slight decrease of total thymocyte numbers at day 21 is likely due to ageing and has to be interpreted accordingly. Further, involution should be considered when comparing other studies with differently aged mice.

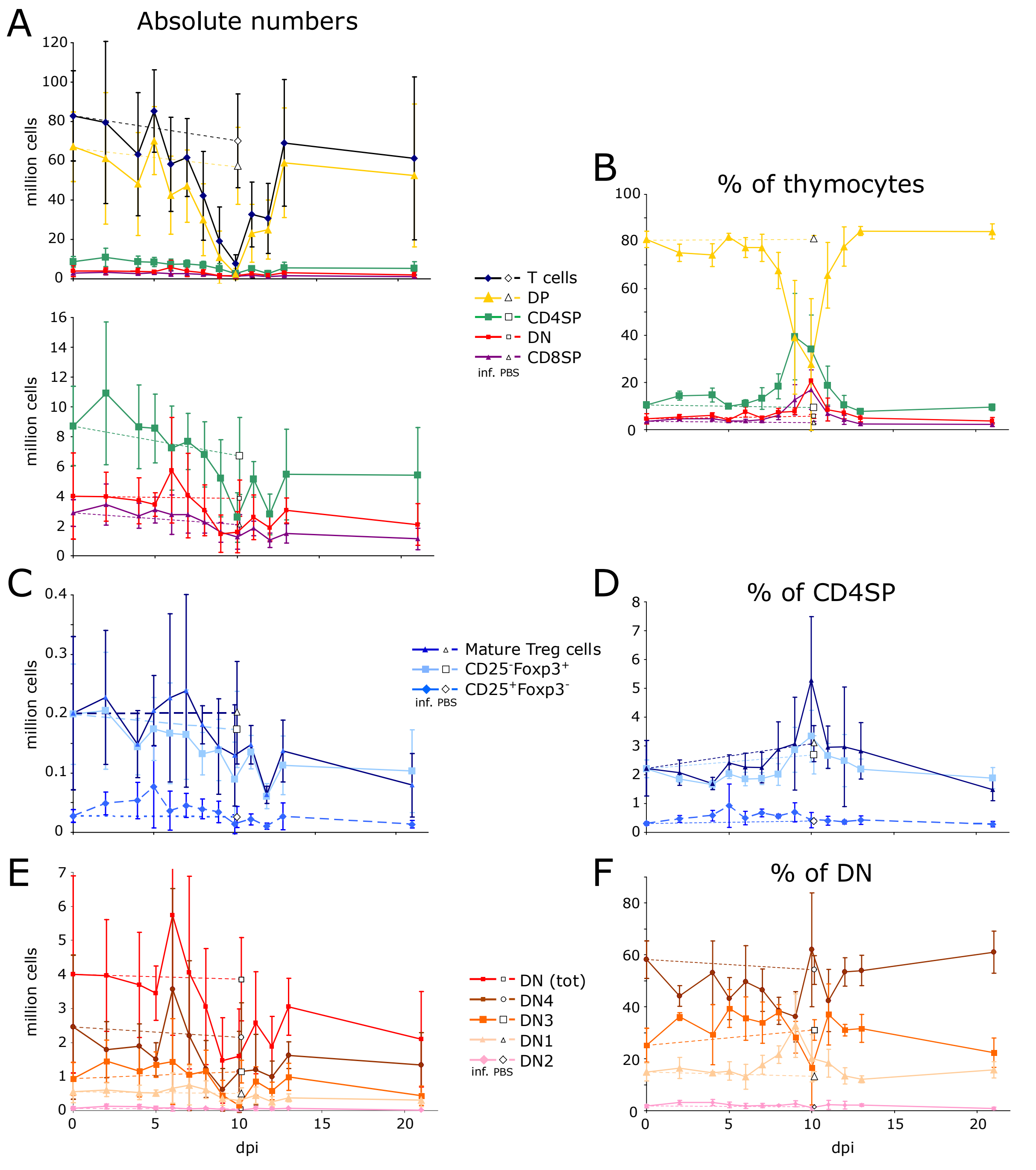

**Figure 'Dynamics of major thymocyte populations upon IAV infection'** Foxp3$^{hCD2}$xRag1$^{GFP}$ double reporter mice were infected with IAV and control mice received PBS. Thymi were analyzed at different days post infection (dpi). (A) Absolute numbers of total thymocytes as well as DP, DN, CD4SP and CD8SP thymocytes. (B) Frequencies of indicated thymocyte subsets among total thymocytes. (C+D) Mature Treg cells and Treg cell precursors (CD25$^{+}$Foxp3$^{-}$ and CD25$^{-}$Foxp3$^{+}$) as absolute numbers (C) and frequency of CD4SP thymocytes (D). (E+F) DN thymocyte subsets as numbers (E) and frequencies of total DN thymocytes (F). To monitor the effect of aging during the experiment, all mice are infected at 6 to 7 weeks of age, and a PBS control is added in the curves at day 7.

**Dynamics of newly generated Rag1$^{GFP+}$ and resident and/or recirculating Rag1$^{GFP-}$ cells:** During IAV-induced thymus atrophy, the absolute numbers of CD8SP and total CD4SP thymocytes as well as Treg cells and their precursors seem to be mildly affected, with a reduction around 50 % at the peak (day 10) and a slight decrease maintained after the peak (see Figure ‘Dynamics of major thymocyte populations upon IAV infection’). In order to assess if newly generated Rag1$^{GFP+}$ or resident and/or recirculating Rag1$^{GFP-}$ cells were preferentially affected, each thymocyte population was separated into Rag1$^{GFP+}$ and Rag1$^{GFP-}$ cells. Rag1$^{GFP-}$ cells are rather constant in each population, meaning the impact of the infection is rather on newly generated thymocytes. Further, it has to be noted that a substantial fraction of mature Treg cells and CD25$^{-}$Foxp3$^{+}$ precursors are Rag1$^{GFP-}$. Therefore, it is important to separate Rag1$^{GFP+}$ and Rag1$^{GFP-}$ cell populations for a dynamic modelling approach since they follow different dynamics.

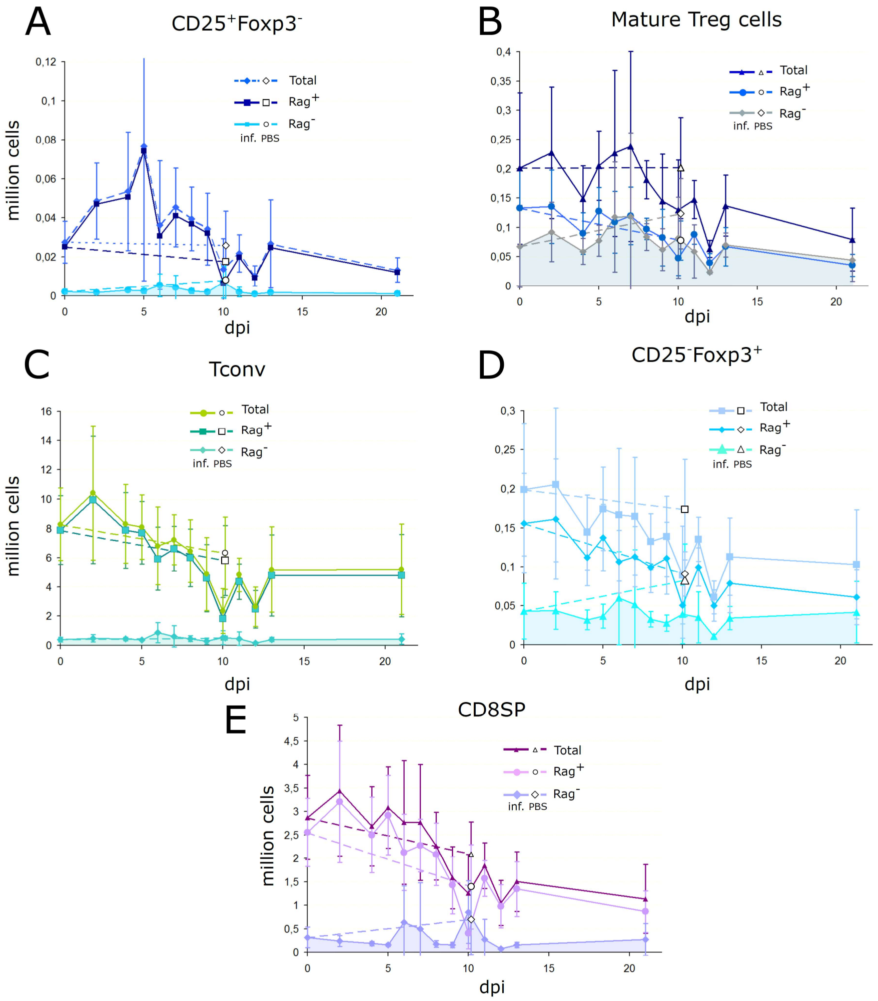

**Figure ‘Dynamics of newly generated Rag1$^{GFP+}$ as compared to Rag1$^{GFP-}$ cells’** Foxp3$^{hCD2}$xRag1$^{GFP}$ double reporter mice were infected with IAV and control mice received PBS. Thymi were analyzed at different days post infection (dpi) and absolute numbers of total as well as Rag1$^{GFP+}$ and Rag1$^{GFP-}$ fractions of indicated thymocyte populations were determined.

To demonstrate that resident and/or recirculating Rag1$^{GFP-}$ cells are not impacted during IAV-induced thymus atrophy, the total amount of Rag1$^{GFP-}$ cells was determined at indicated time points (see Figure 'Dynamics and composition of the pools of Rag1GFP- cells during IAV-induced thymus atrophy'). At the SP thymocyte stage, the total amount of Rag1$^{GFP-}$ cells are rather stable (A), although subject to variation, while the Rag1$^{GFP+}$ population of Treg cell precursors are more impacted than the Rag1$^{GFP-}$ populations (B+D). The composition of the Rag1$^{GFP-}$ pool of SP thymocyte subsets is quite unchanged during thymus atrophy for the mature Treg cells and Treg cell precursors (C). Yet, at peak of thymus atrophy, a slight variation is observed for Tconv and CD8SP Rag1$^{GFP-}$ cells. Furthermore, the frequencies of Rag1$^{GFP-}$ cells inside each thymocyte population are slightly increased at the peak of thymus atrophy (E). Still, the pool of Rag1$^{GFP-}$ cells is rather stable and no specific event of modulated recirculation, death or exit of Rag1$^{GFP-}$ cells could be detected, supporting that Rag1$^{GFP-}$ cells can be modelled as constant in the mathematical model.

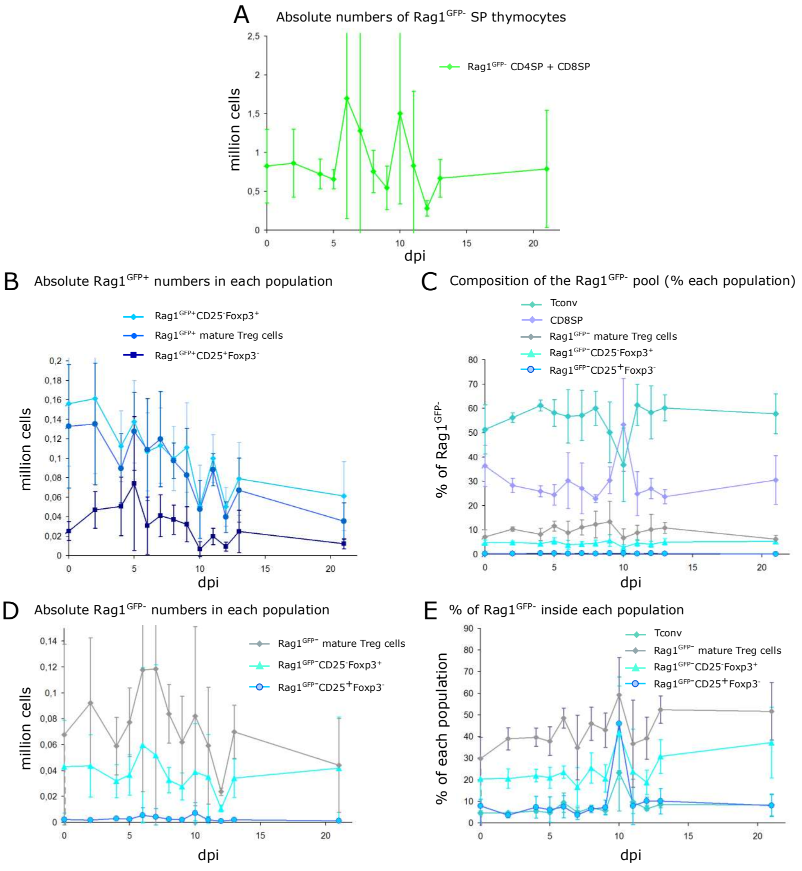

**Figure 'Dynamics and composition of the pools of Rag1$^{GFP-}$ cells during IAV-induced thymus atrophy'** Foxp3$^{hCD2}$xRag1$^{GFP}$ double reporter mice were infected with IAV and control mice received PBS. Thymi were analyzed at different days post infection (dpi) and (A) absolute numbers of Rag1$^{GFP-}$ SP thymocytes, (B+D) total numbers of Rag1$^{GFP+}$ and Rag1$^{GFP-}$ cells within mature Treg cells and their precursors, (C) the composition of the Rag1$^{GFP-}$ compartment and (E) the frequencies of Rag1$^{GFP-}$ cells inside mature Treg cells and their precursors were determined.

**Parameter estimation details:** The non-redundant populations/datasets included in the fitting procedure are: Absolute numbers of DP thymocytes, Rag1$^{GFP+}$ Tconv, Rag1$^{GFP+}$ CD8SP thymocytes, Rag1$^{GFP+}$CD25$^{+}$Foxp3$^{-}$ Treg cell precursors, Rag1$^{GFP+}$CD25$^{-}$Foxp3$^{+}$ Treg cell precursors and Rag1$^{GFP+}$ mature Treg cells; percentages of total (Rag1$^{GFP+}$ and Rag1$^{GFP-}$) DP thymocytes, Tconv and CD8SP thymocytes among total thymocytes; percentages of CD25$^{+}$Foxp3$^{-}$ Treg cell precursors, CD25$^{-}$Foxp3$^{+}$ Treg cell precursors and mature Treg cells among CD4SP thymocytes. The mathematical model presented in the main manuscript only explains the dynamics of newly developing Rag1$^{GFP+}$ cells. However, Rag1$^{GFP-}$ values from the simulations are needed to compare the dataset in percent (that include Rag1$^{GFP-}$ cells). As we saw that Rag1$^{GFP-}$ populations are not strongly impacted by IAV-induced thymus atrophy, they were not simulated but taken constant as their steady-state number value (see Table 'Full dataset of this study'), and included in the calculation of percentages. Finally, we did not simulate the dynamics of DN thymocytes, as the data did not show a clear picture of IAV-induced thymus atrophy, but rather display a high variation. Thus, DN thymocytes were considered constant. In order to calculate percentages of populations among total thymocytes, the experimental number of DN thymocytes was included in the calculation of percentages at each time-point. The open source code for the three model structures was deposited at https://gitlab.com/Moonfit/Balthyse together with the Moonfit program [5], such that simulations and fittings can be reproduced more easily.

The following parameter estimation algorithm was used: Stochastic Ranking Evolutionary Strategy, meaning each individual in the population carries not only a parameter set, but also a mutation speed on its own for each parameter. The SBX cross-over was used, sampling of parents was made proportional to their fitness, defined as the competitive advantage to the worst individual of the population. Offspring did not replace the parents, but only the best individuals were kept to maintain the population size. Mutation followed a normal distribution sampled for all the parameters independently at the same time. Each optimization was performed ten times on a population of 500 individuals along 1,000 generations, as we observed convergence around 300 to 500 generations. Every generation, 20 % of the population was generated as new individuals by cross-over, while 50 % of the population size was generated by mutation. Selection of the best decided which individuals survive (among parents and progeny altogether).

***(next page)*** **Table 'Full dataset of this study'** Absolute numbers and frequencies of all major thymocyte populations measured in this study, as the average/standard deviation for each dpi.

### 1 - Frequencies (average and standard deviation)

| | Population | Alive cells (LD-) | DN1 | DN2 | DN3 | DN4 | DN | DP | CD8SP | | | | Tconvs: CD4+CD8-CD25-Foxp3(huCD2)- | | | CD25-Foxp3(huCD2)+ Precursor | | | CD25+Foxp3(huCD2)- Precursor | | | Mature Treg cells (CD25+Foxp3(huCD2)+) | | |
|---|---|---|---|---|---|---|---|---|---|---|---|---|---|---|---|---|---|---|---|---|---|---|---|---|
| | Definition | | CD44+CD25- | CD44+CD25+ | CD44-CD25+ | CD44-CD25- | Total | | Total | Rag+ | Rag- | CD4SP | Total | Rag+ | Rag- | Total | Rag+ | Rag- | Total | Rag+ | Rag- | Total | Rag+ | Rag- |
| Time (days) | | % of all cells | | | % of DN Total | | | % of alive cells | | % of CD8SP | | % of alive cells | % of CD4SP | % of Tconv | | % of CD4SP | % of Tconv | | % of CD4SP | % of Tconvs | | % of CD4SP | % of Tconv | |
| | | | pctDN1 | pctDN2 | pctDN3 | pctDN4 | pctDN | pctDP | pctSP8 | | pctSP8RagN | pctSP4 | pctTconvtot | | pctTconvRagN | pctTregP2tot | | pctTregP2RagN | pctTregP1tot | | pctTregP1RagN | pctTregtot | | pctTregRagN |
| 0 | | 99.62 | 14.92 | 1.740 | 25.18 | 58.18 | 4.618 | 80.94 | 3.452 | 89.620 | 10.38 | 10.46 | 95.24 | 95.34 | 4.674 | 2.210 | 79.60 | 20.40 | 0.3100 | 92.12 | 7.862 | 2.222 | 70.22 | 29.78 |
| 2 | | 98.90 | 16.44 | 3.070 | 36.26 | 44.20 | 5.354 | 75.18 | 4.642 | 93.140 | 6.85 | 14.42 | 95.60 | 95.44 | 4.550 | 1.854 | 79.46 | 20.54 | 0.4720 | 96.42 | 3.594 | 2.076 | 61.06 | 38.94 |
| 4 | | 97.68 | 14.48 | 3.130 | 29.26 | 53.10 | 6.100 | 74.22 | 4.574 | 92.880 | 7.12 | 14.68 | 96.04 | 94.56 | 5.446 | 1.664 | 78.06 | 21.94 | 0.5840 | 92.72 | 7.272 | 1.698 | 60.40 | 39.60 |
| 5 | | 98.95 | 15.30 | 2.248 | 39.35 | 43.13 | 4.105 | 82.05 | 3.603 | 94.650 | 5.38 | 9.95 | 94.65 | 95.20 | 4.810 | 2.025 | 79.08 | 20.93 | 0.9225 | 93.88 | 6.145 | 2.408 | 62.15 | 37.85 |
| 6 | | 91.22 | 13.04 | 1.836 | 35.50 | 49.60 | 7.542 | 77.38 | 3.600 | 90.260 | 9.75 | 11.05 | 95.40 | 90.62 | 9.378 | 1.854 | 76.46 | 23.54 | 0.5040 | 92.48 | 7.512 | 2.272 | 51.48 | 48.52 |
| 7 | | 98.33 | 17.70 | 1.930 | 33.90 | 46.45 | 4.908 | 77.30 | 4.218 | 94.950 | 5.04 | 13.18 | 95.20 | 95.03 | 4.957 | 1.862 | 83.35 | 16.64 | 0.6767 | 96.30 | 3.705 | 2.255 | 65.12 | 34.88 |
| 8 | | 99.12 | 21.60 | 2.035 | 37.87 | 38.48 | 7.380 | 67.67 | 6.115 | 92.367 | 7.63 | 18.33 | 94.53 | 93.00 | 7.015 | 2.008 | 74.60 | 25.40 | 0.5650 | 93.60 | 6.407 | 2.883 | 54.10 | 45.90 |
| 9 | | 92.89 | 32.76 | 2.687 | 28.31 | 36.21 | 7.801 | 39.20 | 12.643 | 89.629 | 10.36 | 39.53 | 93.34 | 94.31 | 5.680 | 2.861 | 79.40 | 20.60 | 0.6986 | 92.84 | 7.164 | 3.079 | 57.06 | 42.94 |
| 10 | | 96.96 | 20.17 | 1.239 | 16.62 | 61.97 | 20.717 | 27.70 | 16.977 | 39.258 | 60.74 | 34.12 | 90.97 | 76.72 | 23.274 | 3.342 | 58.18 | 41.82 | 0.4311 | 54.08 | 45.922 | 5.277 | 40.86 | 59.14 |
| 11 | | 98.67 | 18.28 | 2.235 | 37.07 | 42.42 | 8.535 | 65.65 | 6.770 | 87.500 | 12.49 | 18.75 | 93.97 | 90.27 | 9.743 | 2.665 | 76.13 | 23.86 | 0.4100 | 92.15 | 7.843 | 2.955 | 63.37 | 36.63 |
| 12 | | 97.24 | 13.39 | 2.208 | 31.04 | 53.40 | 7.138 | 77.82 | 4.148 | 92.300 | 7.72 | 10.54 | 94.18 | 93.36 | 6.610 | 2.492 | 81.22 | 18.78 | 0.3620 | 89.92 | 10.082 | 2.976 | 60.84 | 39.16 |
| 13 | | 98.30 | 12.10 | 2.226 | 31.68 | 53.98 | 4.942 | 84.46 | 2.302 | 89.280 | 10.71 | 7.78 | 94.54 | 91.50 | 8.490 | 2.194 | 69.28 | 30.72 | 0.4300 | 89.92 | 10.082 | 2.830 | 47.58 | 52.42 |
| 21 | | 98.97 | 15.77 | 0.848 | 22.30 | 61.10 | 3.745 | 84.23 | 2.187 | 79.483 | 20.53 | 9.47 | 96.35 | 91.67 | 8.327 | 1.880 | 62.82 | 37.18 | 0.2867 | 91.95 | 8.053 | 1.492 | 48.33 | 51.67 |
| PBS (d10) | | 99.32 | 13.30 | 1.415 | 31.07 | 54.22 | 5.625 | 81.30 | 3.023 | 67.063 | 32.93 | 9.46 | 93.83 | 91.42 | 8.573 | 2.692 | 54.35 | 45.65 | 0.3900 | 75.02 | 24.983 | 3.087 | 39.60 | 60.40 |
| PBS(d21) | | 99.04 | 14.56 | 0.864 | 24.2 | 60.36 | 5.798 | 81.58 | 2.822 | 74.1 | 25.9 | 9.404 | 94.18 | 89.02 | 10.99 | 2.792 | 50.30 | 49.70 | 0.276 | 82.04 | 17.956 | 2.77 | 39.78 | 60.22 |

**Standard Deviations**

| | | | DN1 | DN2 | DN3 | DN4 | DN | DP | CD8SP | | | | Tconvs: CD4+CD8-CD25-Foxp3(huCD2)- | | | CD25-Foxp3(huCD2)+ Precursor | | | CD25+Foxp3(huCD2)- Precursor | | | Mature Treg cells (CD25+Foxp3(huCD2)+) | | |
|---|---|---|---|---|---|---|---|---|---|---|---|---|---|---|---|---|---|---|---|---|---|---|---|---|
| | | % of all cells | CD44+CD25- | CD44+CD25+ | CD44-CD25+ | CD44-CD25- | Total | | Total | Rag+ | Rag- | CD4SP | Total | Rag+ | Rag- | Total | Rag+ | Rag- | Total | Rag+ | Rag- | Total | Rag+ | Rag- |
| 0 | | 0.045 | 3.429 | 0.511 | 6.540 | 7.244 | 2.263 | 3.480 | 0.517 | 4.867 | 4.872 | 1.347 | 1.110 | 0.956 | 0.941 | 0.309 | 9.603 | 9.603 | 0.056 | 3.072 | 3.090 | 0.962 | 9.857 | 9.857 |
| 2 | | 0.418 | 4.083 | 0.826 | 1.498 | 4.007 | 1.039 | 3.681 | 0.791 | 1.508 | 1.534 | 1.866 | 0.430 | 0.832 | 0.847 | 0.128 | 4.497 | 4.497 | 0.115 | 1.213 | 1.237 | 0.438 | 5.114 | 5.114 |
| 4 | | 0.870 | 0.576 | 1.152 | 11.567 | 12.295 | 1.134 | 4.888 | 0.978 | 1.219 | 1.216 | 2.954 | 0.439 | 0.757 | 0.779 | 0.146 | 2.893 | 2.893 | 0.187 | 5.046 | 5.050 | 0.216 | 4.235 | 4.235 |
| 5 | | 0.755 | 3.595 | 1.125 | 7.230 | 8.120 | 0.692 | 1.466 | 0.405 | 1.380 | 1.389 | 0.592 | 0.827 | 0.535 | 0.537 | 0.151 | 4.789 | 4.789 | 0.749 | 5.932 | 5.918 | 0.286 | 6.556 | 6.556 |
| 6 | | 7.672 | 4.747 | 0.989 | 8.385 | 13.898 | 2.129 | 4.303 | 0.748 | 4.196 | 4.189 | 1.574 | 0.608 | 4.458 | 4.463 | 0.233 | 2.808 | 2.808 | 0.235 | 5.129 | 5.139 | 0.479 | 4.624 | 4.624 |
| 7 | | 1.354 | 3.660 | 0.818 | 7.944 | 8.153 | 0.852 | 5.876 | 1.241 | 2.760 | 2.741 | 4.606 | 0.657 | 1.958 | 1.972 | 0.207 | 8.879 | 8.886 | 0.129 | 2.065 | 2.062 | 0.529 | 15.011 | 15.011 |
| 8 | | 0.306 | 3.356 | 0.232 | 3.022 | 5.232 | 1.679 | 7.702 | 1.823 | 1.172 | 1.181 | 5.328 | 1.148 | 1.176 | 1.199 | 0.374 | 6.960 | 6.960 | 0.062 | 1.631 | 1.648 | 0.953 | 7.649 | 7.649 |
| 9 | | 9.952 | 12.313 | 1.200 | 6.165 | 9.683 | 1.384 | 24.189 | 6.439 | 2.835 | 2.850 | 18.394 | 2.484 | 1.643 | 1.659 | 0.780 | 6.272 | 6.272 | 0.325 | 3.429 | 3.424 | 1.610 | 7.978 | 7.978 |
| 10 | | 1.108 | 9.157 | 1.019 | 14.428 | 21.871 | 14.248 | 28.002 | 8.410 | 28.633 | 28.636 | 14.640 | 2.944 | 17.839 | 17.842 | 0.897 | 21.478 | 21.478 | 0.264 | 21.636 | 21.636 | 2.213 | 17.393 | 17.393 |
| 11 | | 0.532 | 5.452 | 1.789 | 11.779 | 12.046 | 4.847 | 13.949 | 3.114 | 17.168 | 17.174 | 8.326 | 1.226 | 10.473 | 10.467 | 0.429 | 19.515 | 19.518 | 0.154 | 8.641 | 8.649 | 0.740 | 20.144 | 20.144 |
| 12 | | 1.717 | 3.202 | 1.469 | 2.255 | 5.443 | 2.627 | 8.356 | 1.851 | 2.355 | 2.359 | 3.931 | 2.905 | 0.999 | 0.989 | 0.902 | 5.630 | 5.630 | 0.064 | 1.867 | 1.860 | 2.072 | 9.716 | 9.716 |
| 13 | | 0.886 | 1.306 | 0.494 | 5.413 | 5.894 | 1.558 | 1.993 | 0.580 | 1.516 | 1.525 | 1.050 | 1.176 | 1.513 | 1.484 | 0.348 | 7.841 | 7.841 | 0.149 | 5.877 | 5.869 | 0.983 | 6.295 | 6.295 |
| 21 | | 0.700 | 3.175 | 0.492 | 5.635 | 8.092 | 1.444 | 3.205 | 0.831 | 15.285 | 15.278 | 1.317 | 0.799 | 5.001 | 5.008 | 0.366 | 16.306 | 16.306 | 0.091 | 5.107 | 5.105 | 0.390 | 13.301 | 13.301 |
| PBS (d10) | | 0.050 | 1.791 | 0.146 | 4.001 | 5.576 | 0.865 | 1.308 | 0.420 | 7.278 | 7.287 | 0.432 | 1.322 | 2.704 | 2.696 | 0.657 | 9.579 | 9.579 | 0.059 | 5.392 | 5.392 | 0.625 | 5.813 | 5.813 |
| PBS (d21) | | 0.707 | 1.131 | 0.148 | 1.838 | 3.111 | 0.035 | 0.424 | 0.085 | 7.990 | 7.990 | 0.410 | 0.566 | 1.485 | 1.442 | 0.007 | 8.132 | 8.132 | 0.177 | 7.990 | 8.004 | 0.361 | 9.263 | 9.263 |

### 2 - Absolute numbers (average and standard deviation)

| | Population | Alive cells (LD-) | DN1 | DN2 | DN3 | DN4 | DN | DP | CD8SP | | | | Tconvs: CD4+CD8-CD25-Foxp3(huCD2)- | | | CD25-Foxp3(huCD2)+ Precursor | | | CD25+Foxp3(huCD2)- Precursor | | | Mature Treg cells (CD25+Foxp3(huCD2)+) | | |
|---|---|---|---|---|---|---|---|---|---|---|---|---|---|---|---|---|---|---|---|---|---|---|---|---|
| Time (days) | Alive | t cells | CD44+CD25- | CD44+CD25+ | CD44-CD25+ | CD44-CD25- | Total | | Total | Rag+ | Rag- | CD4SP | Total | Rag+ | Rag- | Total | Rag+ | Rag- | Total | Rag+ | Rag- | Total | Rag+ | Rag- |
| Model Name | | ttotal | tDN1 | tDN2 | tDN3 | tDN4 | tDN | tDP | tSP8tot | tSP8P | tSP8RagN | tSP4tot | tTconvtot | tTconvP | tTconvRagN | tTRegP2tot | tTregP2P | tTregP2RagN | tTRegP1tot | tTregP1P | tTregP1RagN | tTregtot | tTregP | tTregRagN |
| 0 | 83065200 | 82757854 | 551435 | 68149 | 935109 | 2452571 | 4007264 | 67168932 | 2870602 | 2552949 | 317654 | 8711057 | 8283783 | 7889550 | 394233 | 198951 | 155876 | 43075 | 27499 | 25238 | 2261 | 200695 | 132949 | 67745 |
| 2 | 79751200 | 79480854 | 609635 | 122305 | 1447842 | 1786955 | 3966737 | 61161025 | 3440560 | 3202513 | 238047 | 10912533 | 10431103 | 9943283 | 487820 | 205043 | 161337 | 43707 | 48674 | 46953 | 1721 | 227431 | 135290 | 92141 |
| 4 | 63471000 | 63255524 | 533107 | 116479 | 1157532 | 1885509 | 3692627 | 48216273 | 2685479 | 2498819 | 186660 | 8661145 | 8314597 | 7870209 | 444387 | 144134 | 112246 | 31888 | 53544 | 50630 | 2914 | 148653 | 89673 | 58979 |
| 5 | 85570500 | 85294876 | 510507 | 78916 | 1356442 | 1504088 | 3450962 | 70219661 | 3079850 | 2920086 | 159764 | 8544403 | 8088385 | 7707806 | 380579 | 174238 | 137503 | 36735 | 76653 | 74120 | 2533 | 205109 | 127714 | 77395 |
| 6 | 58476375 | 58259283 | 662003 | 76205 | 1437192 | 3568510 | 5743911 | 42511414 | 2765285 | 2127906 | 637379 | 7238674 | 6809097 | 5930706 | 878391 | 166592 | 106727 | 59865 | 36320 | 30830 | 5490 | 226564 | 108889 | 117675 |
| 7 | 62151667 | 61648940 | 745112 | 57350 | 1050524 | 2201296 | 4054283 | 47151868 | 2766754 | 2274362 | 492392 | 7676036 | 7227387 | 6613518 | 613869 | 164764 | 113007 | 51757 | 45359 | 41041 | 4318 | 238346 | 119809 | 118537 |
| 8 | 42387000 | 42194267 | 620631 | 60351 | 1152253 | 1222101 | 3055336 | 30070431 | 2260535 | 2086557 | 173978 | 6807965 | 6454650 | 5992136 | 462514 | 132447 | 99843 | 32604 | 39465 | 36892 | 2573 | 181254 | 97375 | 83879 |
| 9 | 19320429 | 19160131 | 364422 | 43393 | 453663 | 611438 | 1472915 | 10897267 | 1583801 | 1427827 | 155975 | 5206147 | 4888274 | 4592237 | 296037 | 138850 | 111087 | 27762 | 34242 | 32187 | 2054 | 144509 | 82548 | 61961 |
| 10 | 7901000 | 7864765 | 290689 | 12746 | 149573 | 1141772 | 1594780 | 2427653 | 1257624 | 404142 | 853482 | 2584708 | 2351707 | 1827591 | 524116 | 89488 | 50534 | 38954 | 13742 | 6534 | 7208 | 129684 | 47572 | 82113 |
| 11 | 32765000 | 32666686 | 447926 | 59733 | 866809 | 1202589 | 2577057 | 23107147 | 1845894 | 1569909 | 275985 | 5136588 | 4832164 | 4372078 | 460086 | 135141 | 99904 | 35237 | 21830 | 19795 | 2035 | 147384 | 88210 | 59173 |
| 12 | 30828000 | 30718141 | 262150 | 48264 | 579378 | 985478 | 1875270 | 25008661 | 1051690 | 978144 | 73546 | 2782520 | 2648001 | 2476849 | 171152 | 60839 | 50122 | 10717 | 10337 | 9303 | 1034 | 63272 | 39510 | 23762 |
| 13 | 69442200 | 69024577 | 371619 | 65688 | 988981 | 1626656 | 3052944 | 59001651 | 1501760 | 1344431 | 157329 | 5468222 | 5191264 | 4784013 | 407251 | 113024 | 78910 | 34114 | 26691 | 24824 | 1867 | 137158 | 67193 | 69965 |
| 21 | 61371000 | 61186305 | 306618 | 15756 | 437437 | 1340542 | 2100353 | 52545488 | 1135870 | 865114 | 270756 | 5404594 | 5208662 | 4776181 | 432481 | 103077 | 61198 | 41880 | 13359 | 12096 | 1263 | 79462 | 35389 | 44074 |
| PBS (d10) | 70628667 | 70151033 | 504365 | 54355 | 1146426 | 2150643 | 3855789 | 57491726 | 2092278 | 1393744 | 698534 | 6711239 | 6310362 | 5805031 | 505331 | 173424 | 90767 | 82656 | 25931 | 17733 | 8198 | 201422 | 77888 | 123534 |
| PBS(d21) | 99298714 | 98828065 | 709628 | 46941 | 1324465 | 2789272 | 4870305 | 81982183 | 2454076 | 1954098 | 499978 | 9521500 | 9045608 | 8172386 | 873222 | 212830 | 115561 | 97270 | 25988 | 21624 | 4364 | 236893 | 94093 | 142801 |

**Standard Deviations**

| | | | DN1 | DN2 | DN3 | DN4 | DN | DP | CD8SP | | | | Tconvs: CD4+CD8-CD25-Foxp3(huCD2)- | | | CD25-Foxp3(huCD2)+ Precursor | | | CD25+Foxp3(huCD2)- Precursor | | | Mature Treg cells (CD25+Foxp3(huCD2)+) | | |
|---|---|---|---|---|---|---|---|---|---|---|---|---|---|---|---|---|---|---|---|---|---|---|---|---|
| | | | CD44+CD25- | CD44+CD25+ | CD44-CD25+ | CD44-CD25- | Total | | Total | Rag+ | Rag- | CD4SP | Total | Rag+ | Rag- | Total | Rag+ | Rag- | Total | Rag+ | Rag- | Total | Rag+ | Rag- |
| 0 | 22956557 | | 314785 | 44067 | 480388 | 2118932 | 2895979 | 17662302 | 897745 | 730066 | 217113 | 2661534 | 2475134 | 2334784 | 165293 | 84696 | 63858 | 35863 | 10722 | 9782 | 1344 | 129022 | 63463 | 69842 |
| 2 | 41231069 | | 198522 | 65923 | 623775 | 828291 | 1642017 | 33393734 | 1392645 | 1295186 | 116687 | 4782949 | 4574633 | 4344473 | 247307 | 98501 | 76897 | 24093 | 19441 | 18784 | 811 | 112680 | 62539 | 50447 |
| 4 | 31263417 | | 211865 | 59350 | 762245 | 660211 | 1541667 | 26110401 | 847143 | 804641 | 48004 | 2822604 | 2694625 | 2586437 | 120192 | 47707 | 36360 | 12773 | 30394 | 30161 | 608 | 56180 | 35540 | 22219 |
| 5 | 20918783 | | 88105 | 48030 | 373776 | 485485 | 797935 | 17182454 | 867647 | 846533 | 34384 | 2318498 | 2189120 | 2123808 | 68263 | 53121 | 42457 | 15553 | 68989 | 68688 | 1345 | 59052 | 39968 | 26512 |
| 6 | 23976430 | | 440045 | 88230 | 1260169 | 2960016 | 3536231 | 19852114 | 1318771 | 812908 | 779609 | 2822800 | 2667302 | 2161217 | 694025 | 84997 | 40158 | 59733 | 32941 | 29400 | 5847 | 141700 | 52521 | 93884 |
| 7 | 19737845 | | 580141 | 19237 | 315628 | 2199514 | 2835250 | 18371472 | 1235634 | 564420 | 987053 | 1871382 | 1703928 | 1522504 | 725766 | 75367 | 38764 | 70025 | 20151 | 21507 | 6236 | 162526 | 48935 | 142768 |
| 8 | 22536690 | | 255354 | 29418 | 628150 | 831854 | 1708255 | 18164289 | 726087 | 664639 | 69869 | 2194395 | 2134018 | 1962239 | 185758 | 35657 | 32904 | 9861 | 16352 | 15156 | 1378 | 32313 | 18167 | 22749 |
| 9 | 17435740 | | 214511 | 40141 | 430604 | 619458 | 1251795 | 13357865 | 661647 | 607911 | 59807 | 2577695 | 2499634 | 2299720 | 211918 | 51713 | 45932 | 10497 | 18370 | 18138 | 765 | 80523 | 48098 | 35568 |
| 10 | 4394643 | | 235791 | 11667 | 149492 | 1187965 | 1384379 | 2927461 | 823510 | 326963 | 667160 | 1659209 | 1536539 | 1444213 | 483698 | 62554 | 42309 | 37508 | 15612 | 7671 | 8123 | 85375 | 29717 | 69122 |
| 11 | 16465836 | | 244907 | 54884 | 421392 | 1039934 | 1497297 | 14713214 | 488756 | 380439 | 427883 | 1191379 | 1157556 | 1221182 | 456236 | 28273 | 24407 | 32784 | 9473 | 8415 | 2460 | 32341 | 16546 | 45100 |
| 12 | 17836533 | | 147032 | 38767 | 261841 | 467316 | 889368 | 15090284 | 479081 | 458572 | 23935 | 1378892 | 1351235 | 1280848 | 75185 | 20994 | 20063 | 2639 | 5119 | 4683 | 496 | 14467 | 15716 | 2871 |
| 13 | 32167383 | | 116792 | 15907 | 400797 | 390216 | 834611 | 27825322 | 636798 | 582534 | 58706 | 3041982 | 2928954 | 2783315 | 154866 | 49425 | 37532 | 14803 | 22487 | 21858 | 788 | 51918 | 32808 | 20514 |
| 21 | 41420894 | | 196380 | 14961 | 277115 | 947699 | 1403399 | 36315660 | 729501 | 443833 | 337156 | 3224242 | 3104820 | 2808592 | 360686 | 69882 | 35276 | 39450 | 6218 | 5005 | 1325 | 53480 | 18718 | 36311 |
| PBS (d10) | 23745851 | | 170423 | 17880 | 333171 | 1021187 | 1222865 | 19623814 | 682900 | 896512 | 749429 | 2593777 | 2476419 | 2397069 | 139268 | 64093 | 28520 | 46582 | 17495 | 7963 | 10711 | 86437 | 33277 | 59762 |
| PBS (d21) | 66636281 | | 421166 | 34050 | 950367 | 1361402 | 2695884 | 55252833 | 1540716 | 1423763 | 320856 | 7181925 | 6895781 | 6308401 | 630434 | 116373 | 79722 | 48298 | 20566 | 17175 | 3933 | 168805 | 69007 | 104276 |

**Table S2:** Absolute numbers and frequencies of all subpopulations masured in the thymus, as the average/standard deviation for each time point of infection.
